# Reproducible differentiation of pure ovarian support cells from clinical-grade hiPSCs as a novel infertility treatment

**DOI:** 10.1101/2024.04.29.591741

**Authors:** Bruna Paulsen, Ferran Barrachina, Alexander D. Noblett, Mark Johnson, Simone Kats, Sabrina Piechota, Maria Marchante, Alexandra B. Figueroa, Kathryn S Potts, Graham Rockwell, Alexa Giovannini, Christian C. Kramme

**Author notes:** Corresponding Author: Christian C. Kramme Gameto Inc., 430 E. 29^th^ St Fl 14 New York, New York 10016.

## Abstract

In vitro maturation (IVM) is an infertility treatment used during in vitro fertilization (IVF) procedures in which immature oocytes are matured outside the body, limiting the excessive hormone doses required for retrieval of ready-to-fertilize oocytes. To overcome the historically low embryo formation rate associated with IVM, we have recently demonstrated that co-culture of hiPSC-derived ovarian support cells (OSCs) yielded higher rates of oocyte maturation and euploid embryo formation, by mimicking the complex ovarian environment in vitro, offering a novel solution to overcome the IVM main limitation. To translate this process into clinics, we sourced and engineered a compliant female clinical-grade (CG) hiPSC line to derive OSCs with similar quality attributes and clinical outcomes to results previously demonstrated with a research hiPSC line. We further optimized our manufacturing protocols to enable increased scale and substituted reagents with appropriate higher-quality alternatives. This strategic approach to product development has successfully met scalable manufacturing needs and ultimately resulted in a product of improved reproducibility, purity, and efficacy. Our findings support the use of a similar strategy to fine-tune hiPSC-derived products facilitating translation to clinical applications.

## Introduction

Assisted reproductive technologies (ART) refer to a range of medical procedures targeting fertility treatments ^1^, with *in vitro* fertilization (IVF) and egg freezing being two of the most commonly used approaches. An additional approach, known as *in vitro* maturation (IVM), relies on driving oocyte maturity *in vitro*, as an alternative for the invasive hormonal injections required to recruit mature oocytes *in vivo*. Despite making ART processes less expensive and painful for patients ^2^, IVM is historically limited by poor outcomes, which has motivated the search for solutions to overcome these challenges to increase access to fertility treatments ^3–5^. The ability to recreate the ovarian microenvironment in a dish offers an opportunity to mimic the dynamic environment necessary to support oocyte maturation *in vitro* and improve the outcomes associated with IVM ^6–8^.

Emerging evidence showcasing the application of human induced pluripotent stem cells (hiPSCs) in wide-ranging cell therapies ^9^ provide a promising solution for IVM. hiPSCs have the potential to be differentiated into any cell type in the body, including ovarian support cells (OSCs) carrying phenotypic and functional similarities to the cells of the endogenous ovary ^10–13^. Recently, we demonstrated that by modulating the expression of key transcription factors involved in granulosa cell specification, differentiation of hiPSCs could be orchestrated with greater accuracy leading to the efficient generation of functionally mature ovarian cell types, known as ovarian support cells (OSCs) ^11^. When co-cultured with human immature oocytes, these OSCs enhance oocyte maturation, as well as euploid blastocyst formation rates by providing a dynamic and more physiologically relevant, ovarian-like environment compared to the IVM standard of care ^7,8^, a method that was named OSC-IVM.

Nevertheless, implementation of this method in clinical practice requires a meticulous approach to address several process challenges, notably regarding scaling up manufacturing to meet market demand and ensuring regulatory compliance, including product quality assurance (identity, reproducibility, and potency), as well as safety for human use. Here, we described our strategic plan to achieve clinical translation, including substitution of raw materials by higher quality alternatives, which highlighted the importance of the matrix in modulating the phenotype of the cells during transcription factor-driven differentiation. Additionally, reproducibility and purity of multiple lots of OSCs was demonstrated through the generation of a comprehensive single-cell RNA sequencing atlas encompassing over 75,000 cells, which also provided insights about the mechanism of action of these cells during OSC-IVM. Finally, we demonstrated the generation of a clinical-grade hiPSC line (CG-hiPSC) with comparable quality attributes and potency to the research-use-only cell line (RUO-hiPSC) used for developing and refining the technology. Altogether, these data outline the systematic approach that successfully resulted in a scalable and controlled manufacturing process, ultimately resulting in a more consistent and functional product ready to be translated into clinical settings.

## Results

### 1. Transcription factor mediated differentiation generates ovarian support cells in different stages of ovarian development and folliculogenesis

Consistency and reproducibility are essential traits to ensure the performance and reliability of a therapeutic in clinical use. Manufacturers of cell-derived therapeutics have an intrinsic challenge to maintain and control an extremely dynamic and plastic system to ensure minimal variability on the final product’s critical attributes, particularly when it relates to function. Therefore a thorough characterization of the cellular outcome of multiple batches, combined with a clear understanding of in-process factors that have the potential to impact cellular phenotypes is imperative to de-risk the manufacturing process. We previously reported a novel method to produce highly functional hiPSC-derived OSCs ^11^ and demonstrated they can be utilized as a co-culture supplement to significantly improve the outcomes of *in vitro* maturation of immature human oocytes ^7^. A product of this nature would benefit a broad range of patients in medical need for ART treatments, in addition to providing a cheaper and patient-friendly alternative for individuals interested in IVF or egg freezing with a lower financial and medical burden. To evaluate the feasibility of utilizing a gene-modified hiPSC line, harboring three inducible transcription factors (*NR5A1*, *RUNX2*, and *GATA4*), as a source to generate consistent and functional OSCs, we compared six independent batches of hiPSC-derived OSCs, all from a research-use-only (RUO)-hiPSC line, produced over eight months by multiple operators following a standard operating procedure (**Table 1**). Differentiation of OSCs mediated by overexpression of inducible transcription factors is a fast and straightforward process compared to standard protocols that rely on small molecules to recapitulate developmental trajectories (**Figure 1a**). After 5 days of induction onto matrigel (M), RUO-hiPSCs multiply 5.63±2.85 times and acquire morphological features that resemble human granulosa cells, such as clusters of cells with spiky edges and granules observed in the cell body (**Figure 1a**). Differentiated RUO-OSC-M also expresses CD82, a well-characterized marker of granulosa cell-fate ^11,14^, indicating successful differentiation into the desired cell type (**Figure 1b**).

**Figure 1:**
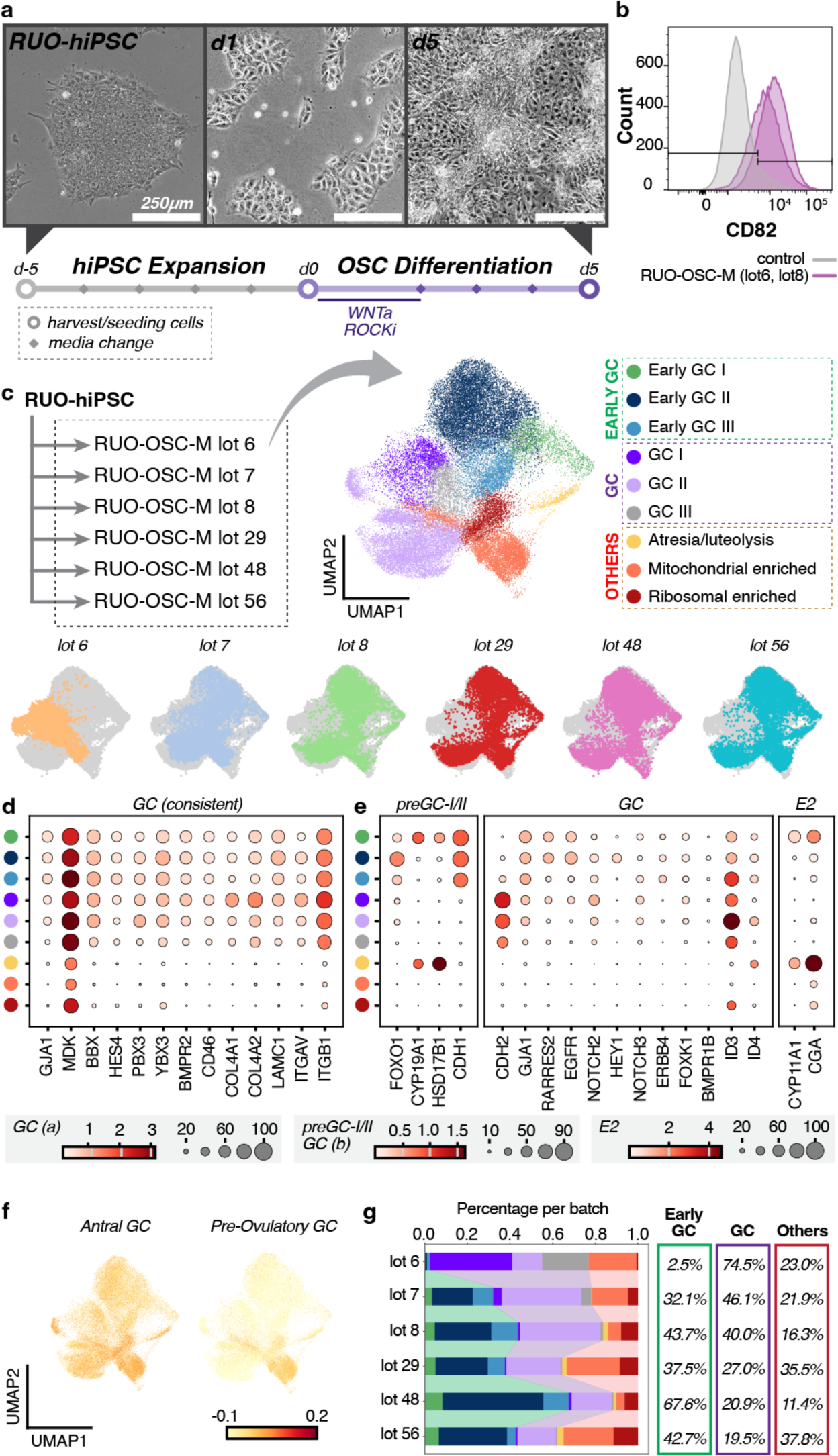
Straightforward and robust differentiation of ovarian support cells (OSCs) derived from a research-use-only (RUO) hiPSC line. a) Timeline and representative images of the RUO-hiPSC expansion followed by differentiation onto matrigel (M) into RUO-OSC-M. Images displayed show the cells on day 5 of hiPSC expansion and days 1 and 5 of OSC differentiation. Scale bar, 250 μm. b) Flow cytometry analysis of the granulosa cell marker, CD82, in the RUO-OSC-M and the negative (unstained) control. d) Dotplot representing the expression of granulosa cell marker genes consistent across GC clusters in the RUO-OSC-M subset. Color cluster legend is found in Figure 1C. Scale represents ‘Mean expression in groups’, ranging from 0 to 3, 0 to 1.5 and 0 to 4, respectively. The circles represent Fraction of cells in the group (%) ranging from 0 to 100, 0 to 90 and 0 to 100, respectively. e) Dotplot representing the expression of preGC-I/II marker genes, granulosa cell marker genes, and steroidogenesis (E2) related genes that supported assignment of each cluster in the RUO-OSC-M subset. Color cluster legend is found in Figure 1C. Scale represents ‘Mean expression in groups’, ranging from 0 to 3, 0 to 1.5 and 0 to 4, respectively. The circles represent Fraction of cells in the group (%) ranging from 0 to 100, 0 to 90 and 0 to 100, respectively. f) UMAP depicting the signature scores for genes corresponding to Antral GC genes and Pre-Ovulatory GC genes. The color scale ranges from -0.1 to 0.2. g) Stacked bar plot depicting the amount of each cluster type found in each lot. The colors correspond to the UMAP cluster colors found in Figure 1C. Overall percentages per group are given to the right of the barplot.

**Table 1:**
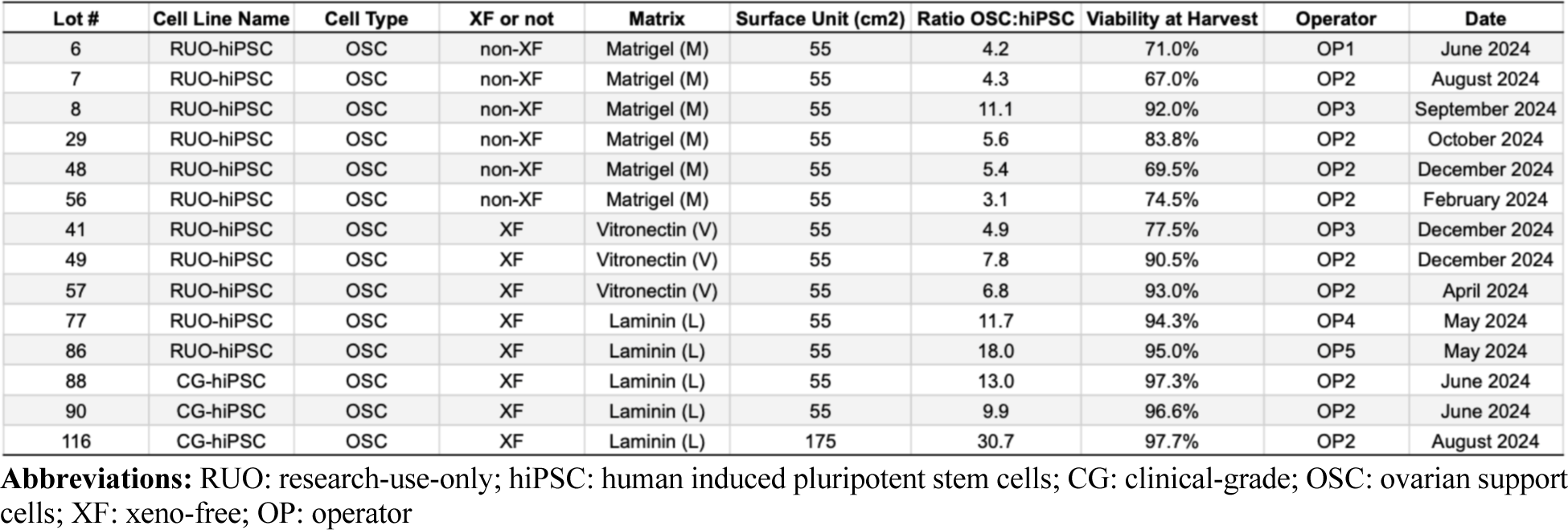
Manifest of Ovarian Support Cell Batch Production and Specifications.

To further characterize the molecular phenotype of the differentiated RUO-OSCs, as well as better understand differences and similarities among independent batches, we performed single cell RNA-sequencing (scRNA-seq) on cryopreserved samples from six batches of differentiation (**Figure 1c**). We identified 15 initial Leiden clusters that were combined by molecular similarities, resulting in nine final clusters. Among the nine final clusters identified, six of them robustly expressed markers previously demonstrated to be differentially expressed in human granulosa cells (*GJA1, MDK, BBX, HES4, PBX3, YBX3, BMPR2, CD46, COL4A1, COL4A2, LAMC1, ITGAV, ITGB1*) compared to other cell types in the developing ovary (**Figure 1d, Supplementary Figure 1**, ^15^. Of note, *MDK,* which encodes the secreted growth factor Midkine, has been shown to improve oocyte maturation and embryo formation in humans and other mammals, corroborating the functional relevance of these cells in the IVM application ^16^. These clusters were all identified as granulosa-like cells, and therefore assigned as the two major classes: ‘Early GCs’ or ‘GCs’. The remaining three clusters were included in a third major class identified as ‘others’, as expression of major markers was not evident in these groups of cells (**Figure 1d**).

Despite the consistent expression of these granulosa cell markers, the class assigned as ‘Early GCs’ also shares transcriptional similarities to preGC-I and -IIa/IIb ^15^, including expression of the genes *FOXO1* and *CDH1* (**Figure 1e**). A subcluster of these cells, labeled as ‘Early GC I’, expresses the aromatase gene *CYP19A1*, which has been described to be upregulated in preGC-Is in the ovarian medulla ^15^, as well as the gene for the chemotactic protein, *RARRES2*, which has been shown to reduce steroidogenesis and block oocyte meiotic progression in bovine models ^17^. The subcluster ‘Early GC II’ also expresses the gene for *RARRES2*, similar to the previous subcluster described, in addition to the receptor *NOTCH2* (**Figure 1e**). It is worth noting that NOTCH signaling pathway is known to be involved in the oocyte-GC crosstalk during folliculogenesis ^14^, and high levels of expression of NOTCH2 and NOTCH3 in cumulus cells have been positively correlated with IVF response ^18^. Finally, in the subcluster ‘Early GC III’ expression of *RARRES2* is no longer observed, as opposed to the previous subcluster; while *NOTCH2* expression continues to be detected at significant levels. Altogether, these patterns of expression suggest that the clusters in the ‘Early GC’ class share transcriptional signature with both granulosa cells and preGC-I and -IIa/IIb, and differential expression of *CYP19A1*, *RARRES2*, and *NOTCH2*, suggest a gradual developmental and functional progression from the subcluster ‘Early GC I’ to ‘Early GC III’.

The class of ‘GCs’ is marked by the expression of *CDH2* in addition to all the other granulosa markers previously described, including the NOTCH2/3 receptors (**Figure 1e**). The subcluster ‘GC I’ is enriched for the genes *NRG1*, *BMPR1B*, and genes of the ERBB family of receptors (**Figure 1e**). *NRG1* has been previously identified to be differentially expressed in preGC-IIa/IIb ^14^ and was found to be expressed and secreted by granulosa cells in response to ovulatory surge ^19^. *BMPR1B*, *EGFR* (*ERBB1*), and *ERBB4* are all receptors previously identified in granulosa cells and known to have counterpart ligands expressed in oocytes (*BMP6*, *TGFA*, and *NRG4*, respectively). These interactions have been proposed to mediate follicular assembly ^14^. The subcluster labeled as ‘GC II’ is enriched by expression of the gene *ID3*, which is a target of the receptor *BMPR2*, also expressed by these cells. Interestingly, although *BMPR2* is expressed by all ‘Early GCs’ and ‘GC’ clusters, the ‘GC II’ is the subcluster with the strongest enrichment of this target gene (**Figure 1e**), suggesting activation of the receptor BMPR2 in these cells. The last subcluster from the ‘GC’ class, ‘GC III’, is composed of cells expressing both *CDH2* and *NOTCH2*, but is not enriched for any of the other genes previously described in the subclusters for this class. In summary, we believe that the three subclusters of ‘GCs’ represent ovarian support cells in slightly different cell states mediated by a distinct combination of active signaling pathways.

The last three subclusters identified (Atresia/luteolysis; Mitochondrial enriched; and Ribosomal enriched) were incorporated into a third class labeled as ‘others’. These clusters have overall lower expression of most markers, including *GJA1* and *CDH2* (**Figure 1d-e**), and reduced levels of these two markers have been previously described in GCs undergoing early stages of atresia ^20^. Interestingly, cells on the ‘Atresia/luteolysis’ subcluster also express genes involved in steroidogenesis, such as *CYP11A1*, *CYP19A1*, and *HSD17B1*, as well as *CGA*, which is an estrogen receptor alpha-responsive gene ^21^. The other two clusters are also enriched for the *CGA* gene, but the top expressed genes in each of the clusters are either mitochondrial genes in the ‘Mitochondrial enriched’ subcluster or ribosomal genes for the ‘Ribosomal enriched’ subcluster. Generally, enrichment of mitochondrial and/or ribosomal genes in scRNA-seq analysis is associated with poor quality cells, further suggesting that these clusters are composed of dying cells. It is important to highlight that it is unclear whether this observation is a consequence of the biology of ovarian support cells or whether it is an artifact of processing and handling the samples.

After identifying that the large majority of the cells in our analysis were classified as granulosa-like cells (‘Early GCs’ and ‘GCs’), we sought to understand whether our protocol gave rise to OSCs in different stages of folliculogenesis or whether cells were overrepresented by a specific follicular stage. For that, we leveraged as a reference a published transcriptome landscape of human folliculogenesis ^14^ to generate gene signature scores that were then applied to our samples. We did not observe a clear representation of either the ‘Primary GC’ or ‘Secondary GC’ stages within our samples, and most of the genes associated with these signature scores were not enriched in the analyzed cells. Conversely, the signature scores for ‘Antral GC’ and ‘Pre-ovulatory GC’ were more clearly represented within the clusters identified in our analysis, and multiple genes driving these signatures seem to be enriched by multiple clusters (**Figure 1f**).

Following the characterization of the cellular outcome resulting from our differentiation process, we investigated how reproducible and consistent independent batches of hiPSC-derived OSCs would be from each other. Overall, all batches analyzed consistently generated clusters from the three major classes previously described (Early GCs, GCs, and others), with five of them very similar in terms of cluster distribution per batch (**Figure 1g**). One sample (RUO-OSC-M lot 6) stood out by the low percentage of ‘Early GCs’ and higher percentage of ‘GCs’ compared to the other batches. This variability may be at least in part attributed to raw materials that are known to have lot-to-lot variability, which can impact the final cellular outcome. During the initial phase of product development, this issue was not a major concern, as a risk assessment of raw materials indicated the possibility of substituting variable reagents with higher quality alternatives in subsequent manufacturing steps, thereby advancing towards meeting clinical standards.

### 2. *In vitro* maturation of human oocytes is robustly achieved by multiple batches of hiPSC-derived ovarian support cells

In view of the variable cellular outcome encountered in the different RUO-OSCs batches analyzed, we sought to verify whether the functional readout of independent batches would be also variable or correlated to the cellular outcome. To assess the functional readout of OSCs, we leveraged results from preclinical studies, including data previously published by our group ^7^. In this study, OSCs were co-cultured with immature cumulus enclosed oocytes obtained from individuals undergoing abbreviated gonadotropin stimulation, and the rate of MII oocyte formation was assessed as a measure of OSC potency (**Figure 2a**). To capture the different spectrum of cell composition variability among the six batches analyzed by scRNA-seq, we performed functional analysis on RUO-OSC-M lot 6, which was overall more represented by ‘GCs’ clusters; RUO-OSC-M lot 8, which contained a balanced representation of ‘Early GCs’ and ‘GCs’ clusters; and RUO-OSC-M lot 56, which was more represented by ‘Early GCs’ and ‘Other’ clusters, with a lower contribution of the ‘GCs’ clusters (**Figure 1g**). Given that the OSCs described herein are intended to be used as a supplement to the conventional media for *in vitro* maturation (IVM), named MediCult IVM Media (Coopersurgical), we utilized this IVM media condition as the control and baseline to assess a positive functional readout triggered by addition of OSCs (**Figure 2a**). For these initial analyses, immature cumulus enclosed oocytes retrieved from each donor were randomly split into two conditions, one composed of conventional IVM Media (Control-IVM group) and another composed of conventionalIVM media supplemented with human OSCs (OSC-IVM group). Following IVM co-culture, MII oocyte formation rate was assessed across groups, demonstrating that all the three individual RUO-OSC-M batches analyzed successfully led to higher MII oocyte formation rate compared to the control group (*p*=0.029, lot 6/control:1.37, lot 8/control: 1.31, lot 56/control: 1.28) (**Figure 2b**). This indicates that despite observing variable cellular outcomes, OSCs robustly retain the ability to improve human oocyte maturation rate compared to control, suggesting that both ‘GCs’ and ‘Early GCs’ may be facilitating oocyte maturation. It is important to emphasize that a larger sample size per batch would be required to understand whether and how cellular composition may differentially impact oocyte maturation.

**Figure 2:**
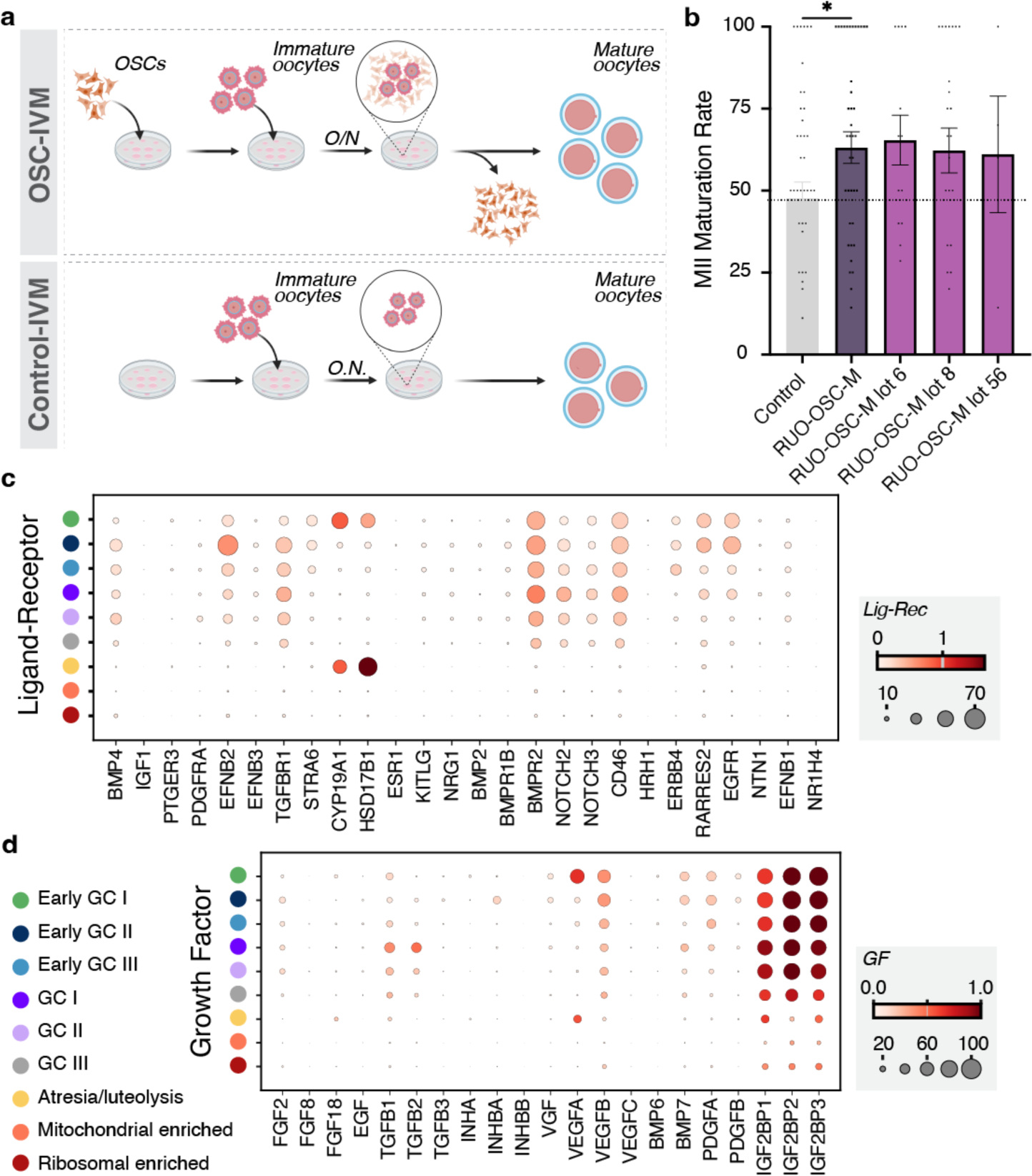
Independent batches of RUO-OSC-M consistently yield successful functional outcomes as evidenced by increased rates of oocyte maturation. a) Schematic representation of the two *in vitro* maturation (IVM) conditions: Control-IVM group containing MediCult-IVM media only, versus OSC-IVM group constituted by MediCult-IVM media supplemented with OSCs. b) Quantification of MII Maturation Rate in Control-IVM (grey) vs OSC-IVM groups. “RUO-OSC-M” displays the combined oocyte maturation rates of 3 separate OSC batches (RUO-OSC-M lots 6, 8, and 56). Data are shown as the mean ± SEM (*p*=0.029, lot 6 vs control:1.37, lot 8 vs control: 1.31, lot 56 vs control: 1.28) c) Dotplot representing the expression of ligand-receptor related genes in the RUO-OSC-M subset in each cluster. Color cluster legend is found in figure 2D. Scale represents ‘Mean expression in groups’, ranging from 0 to 2, and circles represent ‘Fraction of cells in group (%) ranging from 0 to 70. d) Dotplot representing the expression of growth-factor related genes in the RUO-OSC-M subset in each cluster. Scale represents ‘Mean expression in groups’, ranging from 0 to 1, and circles represent ‘Fraction of cells in group (%) ranging from 0 to 100.

To gain insight into the potential mechanism of action associated with these results, we analyzed the expression of key receptors, ligands, and target genes known to have an important role in oocyte and ovarian support cells interactions, on the different classes of cells identified on our samples. Comparison of relative expression of multiple ligand-receptor pairs indicates that ‘GCs’ and ‘Early GCs’ clusters express relative higher levels of *BMP4*, *EFNB2*, *TGFBR1*, *BMPR2*, *NOTCH2*, *NOTCH3*, and *CD46*, suggesting a potential involvement of one of these elements as part of the mechanism of action of these cells (**Figure 2c**). Other receptors, such as *STRA6*, *ERBB4*, *RARRES2*, and *EGFR*, were also detected particularly in the ‘Early GC’ clusters, suggesting that these genes may not be the main drivers of the oocyte maturation process (**Figure 2c**). Additionally, we assessed the expression of growth factors that are also known to be modulated by the interaction between oocytes and somatic cells, and recognized for their involvement in oocyte maturation and folliculogenesis ^14,15^. Among the differentially expressed genes across clusters, *TGFB1* and *TGFB2* show greater enrichment in the GCI cluster; while *VEGFA/B*, as well as *BMP7* and *PDGFA* seem to be all more abundant on the ‘Early GCs’ (**Figure 2d**). All the ‘Early GCs’ and ‘GCs’ clusters were enriched for *IGF2BP1/2/3* (**Figure 2d**). Although these results point to potential pathways involved in the mechanism of action of OSCs responsible for driving oocyte maturation, more studies would be necessary to rule out the involvement of any undetected genes.

### 3. Translation of the protocol towards clinical manufacturing leads to more reproducible cellular outcomes

As part of the strategy to translate the research manufacturing protocol towards clinical standards, we performed a risk assessment on our bill of materials, which led to the substitution of key components of the protocol by higher-quality alternatives, including animal origin-free reagents, GMP-manufactured components, and cell therapy-grade raw materials (**Figure 3a**). We also sought to explore other materials and reagents as factors that might influence OSC differentiation in order to better understand our process and provide a more complete perspective of the significance of the cell culture environment on cell fate determination and specialization. This understanding would drive greater optimization of the process and make the final product more reproducible at the manufacturing stage, ensuring improved efficacy and safety.

**Figure 3:**
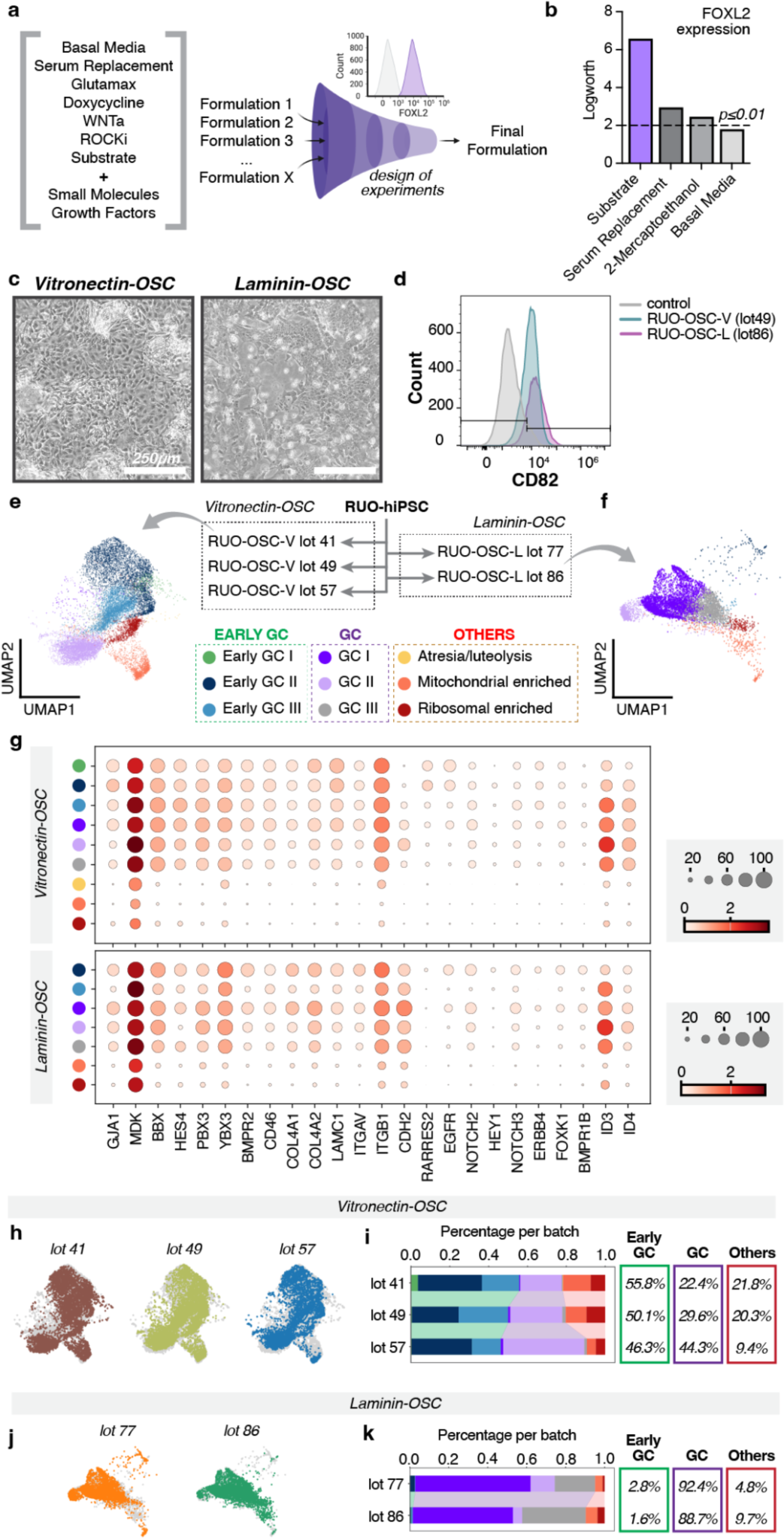
Xeno-free substrate alternatives (vitronectin or laminin) significantly impact OSC differentiation outcomes. a) Schematics of design of experiments (DOE) strategy to optimize manufacturing process. b) Barplot of logworth values of DOE main effect results on FOXL2 expression. Dashed line indicates p≤0.01. c) Images of OSCs in culture on day 5 of differentiation carried out on vitronectin vs laminin-521. Scale bar, 250 μm. d) Flow cytometry analysis of the expression of CD82 in the control, RUO-OSC-V and RUO-OSC-L. e) UMAP projections depicting the RUO-OSC-V subset f) UMAP projections depicting the RUO-OSC-L subset g) Dotplot depicting the expression of granulosa cell markers in the RUO-OSC-V and RUO-OSC-L subsets. Scale represents ‘Mean expression in groups’ ranging from 0 to 3, and the circles represent ‘Fraction of cells in group (%) ranging from 0 to 100. h) UMAP projections depicting each individual lot within theRUO-OSC-V subset. i) Stacked bar plot depicting the amount of each cluster type found in each lot relative to the RUO-OSC-V subset. The colors correspond to the UMAP cluster colors found in figure C. Overall percentages per group are given to the right of the barplot. j) UMAP projections depicting each individual lot within the RUO-OSC-L subset. k) Stacked bar plot depicting the amount of each cluster type found in each lot relative to the RUO-OSC-L subset. The colors correspond to the UMAP cluster colors found in figure C. Overall percentages per group are given to the right of the barplot.

We specifically wanted to explore the combined effects of the inducible transcription factors with media supplements and cell substrates in the growth media environment. Because small molecule-mediated differentiation in supplemented media is an established approach to generate ovarian cell types from hiPSCs, we created a list of potential candidate factors and small molecules from the literature that might influence OSC differentiation from a range of general categories, including basal media, serum replacement, small molecules, and growth factors/morphogens. From the literature sources, we also identified appropriate concentration ranges for each factor (**Supplementary Table 1**). It is understood that the matrix substrate on which cells grow influences a range of factors for hiPSCs in culture, including differentiation, and we added to this list commercially available, animal origin-free extracellular matrix substrates appropriate for translation of clinical cell therapies—laminin and vitronectin ^22^. We included as well different schemes for inducing transcription factor expression, such as doxycycline concentration and doxycycline treatment time (**Supplementary Table 1**). To systematically evaluate the effects of multiple variables at the same time, we employed Design of Experiments (DOE) to create a custom design that included center points for each factor and was optimized for the D-optimality criterion, which is an experimental design matrix that allows us to maximize efficiency and accuracy and minimize uncertainty in the response parameters. For responses in the design, we chose FOXL2 expression and viability, as FOXL2 indicates OSC specification and viability screens for environmental factors that are otherwise unworkable in manufacturing.

We demonstrate that the cell substrate has a clear and strong influence on FOXL2 expression (**Figure 3b**, *p* ≤ 0.01), our desired cellular outcome, far greater than any other factors. This highlights the importance of careful selection of the substrate matrix for hiPSC differentiation. Given that our initial approach utilized Matrigel, known for its batch-to-batch variability, we suggest that the culture platform chosen might have contributed to the observed inconsistency between lots. We also see from the DOE study that doxycycline treatment time is an important factor, but not above the highest significance threshold, while doxycycline concentration is less significant. We established internal controls within this study to include doxycycline-free experimental groups that show a positive correlation between the presence of doxycycline and FOXL2 expression, underscoring the necessity of TF induction and validating our TF-mediated methodology. We are therefore supported in moving forward with our established doxycycline treatment scheme, at least within the parameters of this experimental design. In addition to the cell substrate, two other parameters had significant influence in FOXL2 expression, these being KnockOut Serum and 2-mercaptoethanol. The requirement for serum is expected and, as a result, already included in our process, while supplementation with chemical antioxidants, such as 2-mercaptoethanol, had a clear negative impact on viability that excluded it from practical utility in manufacturing (**Figure 3b**, *p* ≤ 0.01). The other variables in the study were determined to have a relatively lower significance, but considering the dominant effect size of the substrate, there is the potential for effect masking to have occurred, reducing the observable impact of these factors. This result largely met with our expectations however that the TFs play the primary role in driving differentiation independent of the other added small-molecule components. The growth matrix, on the other hand, has a conspicuous influence on directing our engineered hiPSCs towards the targeted OSC phenotype and merits further study and optimization.

Among all the raw materials previously utilized during the generation of OSCs for research purposes, one of the reagents with the highest complexity was Matrigel, a cell substrate derived from Engelbreth-Holm-Swarm mouse sarcoma cells, that contains multiple extracellular matrix components of tissue basement membranes. Due to its nature of production, Matrigel is also subject to significant lot-to-lot variability, which can impact the overall reproducibility of the final product. For substrate alternatives, the most commonly used matrices for hiPSCs cultures are human recombinant laminin and vitronectin. Laminin, in particular, is a primary component of Matrigel, whereas vitronectin contains only trace amounts of matrigel. There are several commercially available laminins and we set out to determine the optimal format for its incorporation into our process. We chose to evaluate different physiologically relevant laminin isoforms, including Laminin-521 and Laminin-511, which are relevant at early stages of development and, therefore, often utilized in stem-cell culture. We seeded our hiPSCs on each matrix and induced differentiation, evaluating cells for FOXL2 expression, as well as metrics of workability, including cost, required concentration, and cell morphology, attachment, and detachment. These results indicate that Laminin-521 best met our targeted outcomes (**Supplementary Figure 2**).

We proceeded by directly comparing differentiation induced onto either human recombinant laminin-521 or vitronectin. All the remaining raw materials were kept the same in both conditions. Initial assessment of cellular morphology during the differentiation process indicated subtle differences between both groups (**Figure 3c**). Vitronectin-OSCs (RUO-OSC-V) presented a larger cell body and organized themselves in more sparse clusters of cells, while laminin-OSC (RUO-OSC-L) generated smaller cells, which organized in compact groups of cells. (**Figure 3c**). Expression of the granulosa-cell marker, CD82 was consistent among these two OSC groups (**Figure 3d**). To better understand the molecular profile of OSCs generated from each of these conditions, as well as understand how reproducible their cellular outcome is, we performed scRNA-seq in three batches of vitronectin-OSC and two batches of laminin-OSCs, and compared them against our previous dataset (**Figure 3e-g, Supplementary Figure 3**). Vitronectin-OSCs are primarily represented by ‘Early GC II’, ‘Early GC III’, and ‘GC III’ (**Figure 3e,g**). In contrast, Laminin-OSCs are mostly distributed throughout the subclusters of the ‘GC’ class, in particular ‘GC I’ and ‘GC III’ (**Figure 3f,g**). Furthermore, the molecular profile of laminin-OSCs closely resembles the cellular outcome observed in the RUO-OSC-M lot 6 batch, which stood out in our previous analysis due to its unique profile among the batches differentiated onto matrigel (**Figure 1c,f**). Notably, OSCs differentiated onto vitronectin had also a higher percentage of cells in the clusters of both ribosomal and mitochondrial gene enrichment (**Figure 3e**). Interestingly, expression of N-cadherin (CDH2), which is a hallmark of the ‘GCs’ subclusters and not present in the ‘Early GCs’, has been described to protect granulosa cells from apoptosis associated with follicular atresia and luteolysis ^23^, and vitronectin (VTN) was identified upregulated in porcine atretic follicles ^24^, suggesting an association of this particular matrix to the higher percentages of cells in the ‘others’ class (Mitochondrial enriched, Ribosomal enriched, atresia/luteolysis).

Despite the differences observed in cellular outcome generated in each of these two conditions, no major differences were noted in cluster distribution among the independent batches generated under the same condition (**Figure 3h-k**), in contrast to the intra-variability observed in the batches differentiated on matrigel (**Figure 1c**). This suggests that changes in the bill of materials, incorporating higher quality reagents, resulted in consistent and reproducible cellular outcomes regardless of the matrix utilized. It is also important to highlight that each independent OSC batch was generated by a different operator, which strengthens the evidence of reproducibility among different lots. Overall, this data indicates that differentiation performed onto laminin-521 and vitronectin generated similar cell fates (‘classes’) to the ones previously characterized, suggesting that overall matrigel-OSC encompasses all likely cellular outcomes. Therefore, as we previously demonstrated that distinct batches of OSCs successfully induce oocyte maturation, it is unlikely that these modifications will impact the function of the cells differentiated onto either laminin or vitronectin. Interestingly, these results demonstrate that the final OSC fate can be modulated by not only the overexpression of the three transcription factors, but can be also significantly influenced by the nature of the matrix utilized as the substrate (**Figure 3h-k**).

### 4. Differentiation over laminin-521 leads to a scalable, pure, and functional population of OSCs

After ensuring that transition to an overall higher quality bill of materials does not compromise reproducibility or lead to a completely novel cellular outcome, we then sought to investigate which of the two conditions (laminin-521 and vitronectin) would yield improved clinical outcome, consequently identifying the condition that should be used for clinical manufacturing. As a measurement of successful clinical outcome, we considered a few parameters that would directly inform the throughput and potency of each condition. For throughput, we compared the ratio of OSC:hiPSC for each batch analyzed per condition (**Table 1**). The condition that allows for a higher yield of harvested viable cells, without changing the initial cell number or surface area for culturing the cells, would make the case for a more scalable alternative. Laminin-OSCs were harvested at 94.63±0.01% viability and during differentiation were multiplied at a ratio of 14.83±4.48 OSC:hiPSCs (**Table 1**). In contrast, vitronectin-OSCs were harvested at 87.00±0.08% viability and were multiplied at a ratio of 6.49±1.43 OSC:hiPSC (**Table 1**).

Most importantly, we must ensure that the OSCs differentiated under these optimized conditions are equally functional (potent) and perform similarly in maturing human oocytes as previously observed (**Figure 2a,b**). With this goal, we co-cultured both laminin-OSCs and vitronectin-OSCs independently with human immature cumulus enclosed oocytes following a similar approach as previously described (see Materials and Methods) and evaluated the MII oocyte maturation rate. We demonstrated that the OSC batches differentiated after bill of material change, namely the OSC-laminin and OSC-vitronectin batches, successfully led to a higher MII formation rate compared to the control (*p*=0.018, RUO-OSC-V lot 41/control:1.08, RUO-OSC-V lot 49/control: 1.36, RUO-OSC-L lot 86/control: 1.27, **Figure 4a,b**). Although both approaches seemed to have generated functional OSCs that contributed to successfully increasing oocyte maturation rate compared to control conditions, the vitronectin-OSC condition resulted in variable functional outputs, being the potency of one of the batches (RUO-OSC-V lot 41) considerably inferior than all the other batches previously analyzed in this study (**Figure 4b**). It is important to note that in this stage of product development, a successful functional outcome informs the potential of both conditions to generate functional OSCs.

**Figure 4:**
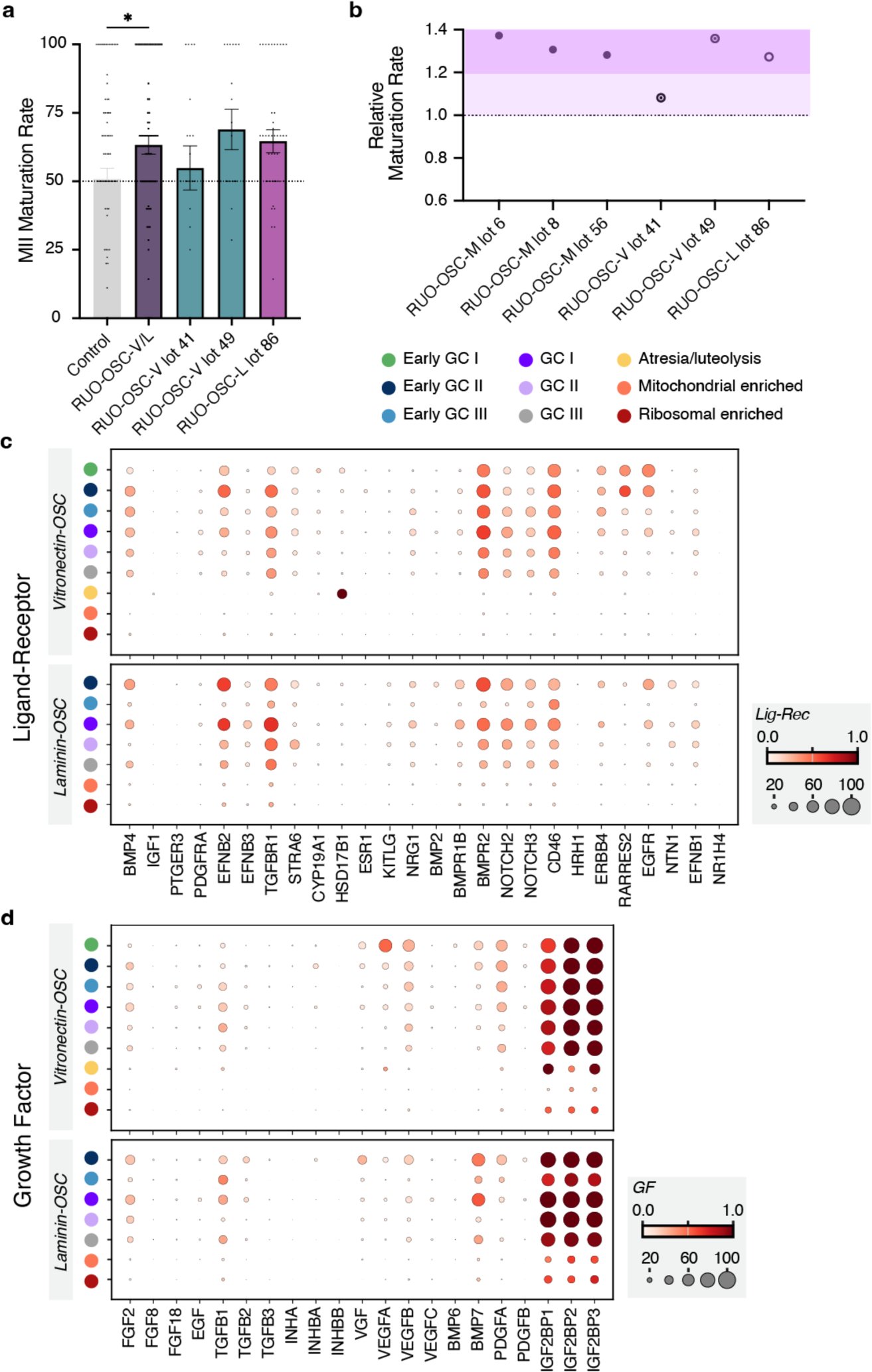
Xeno-free OSC differentiation lead to successful functional outcome measured by higher rates of MII formation. a) Comparison of MII Maturation Rates between control-IVM group (grey) and OSC-IVM groups (RUO-OSC-V/L, RUO-OSC-V lot 41,RUO-OSC-V lot 49, and RUO-OSC-L lot 86). Maturation rates of three separate batches RUO-OSC-V lot 41,RUO-OSC-V lot 49, and RUO-OSC-L lot 86) are combined in the RUO-OSC-L/V bar, with additional bars displaying the individual maturation rates of each batch. Data is the mean ± SEM (*p*=0.018, lot 41/control:1.08, lot 49/control: 1.36, lot 86/control: 1.27) b) Relative MII maturation across OSC batches differentiated on different matrices; matrigel, vitronectin, and laminin 521. c) Dotplot depicting the expression of Ligand-Receptor genes in the RUO-OSC-V and RUO-OSC-L subsets.Scale represents ‘Mean expression in groups’ ranging from 0 to 1, and the circles represent ‘Fraction of cells in group (%) ranging from 0 to 100. d) Dotplot depicting the expression of Growth Factor genes in the RUO-OSC-V and RUO-OSC-L subsets.Scale represents ‘Mean expression in groups’ ranging from 0 to 1, and the circles represent ‘Fraction of cells in group (%) ranging from 0 to 100.

Despite the overall higher rates of MII oocyte maturation in both laminin-OSC and vitronectin-OSC compared to control, it is clear that these two conditions are composed of cells with different phenotypic compositions (**Figure 3**) and, therefore, may drive oocyte maturation through different mechanisms. Hence, to further investigate potential OSC-oocyte interactions and involvement of key signaling pathways associated with ovarian follicle assembly and oocyte meiotic progression, we leveraged published data previously described in endogenous tissue ^14,15^ to characterize how these genes were expressed in OSCs from each condition (**Figure 4c,d**). Overall expression of ligand-receptors pattern was similar between vitronectin-OSC and laminin-OSC, with *BMPR1B* slightly more enriched in the laminin-OSC group compared to the vitronectin-OSC group (**Figure 4c**). This suggests that distinct subgroups of cells (‘Early GCs’ and ‘GCs’) are likely equally receptive to paracrine and/or autocrine signaling. Comparison of expression pattern of growth factors among both groups indicates a few differences (**Figure 4d**). For instance, *VEGFA/B* and *PDGFA* seem to be more enriched in the ‘Early GCs’ and therefore in the vitronectin-OSC samples; while *BMP7* seems to be more expressed in the ‘Early GC II’, ‘GC I’, and ‘GC III’ of the laminin-OSC samples (**Figure 4d**). Interestingly, both *BMP4* and *BMP7* have been identified to differentially regulate FSH-dependent estradiol and progesterone production ^25,26^, suggesting a potential contribution to the OSC-laminin mechanism of action.

### 5. Generation of a clinical-grade hiPSC line with similar attributes to RUO-hiPSC line

Another indispensable modification prior to fully translating this technology into clinical manufacturing is the utilization of clinical- and commercial-grade starting material. Our previous studies were performed with an hiPSC line, considered to be for research-use-only (**RUO-hiPSC**). Sourcing a clinical-grade hiPSC line generated from an allogeneic female donor with proper consent and eligibility (see Materials and Methods) was paramount to enable the progression of these studies (**Figure 5a**). To minimize discrepancies between the results from the original RUO-hiPSC line, which provided the foundation for initial preclinical studies and potency tests, and the clinical-grade (CG)-hiPSC line, we applied the same manufacturing strategy used to engineer the original line ^11^ for the generation of the clinical-grade one with the goal to demonstrate that the same approach can be applied in both contexts to draw qualitative and functional equivalency between the two resulting lines.

**Figure 5:**
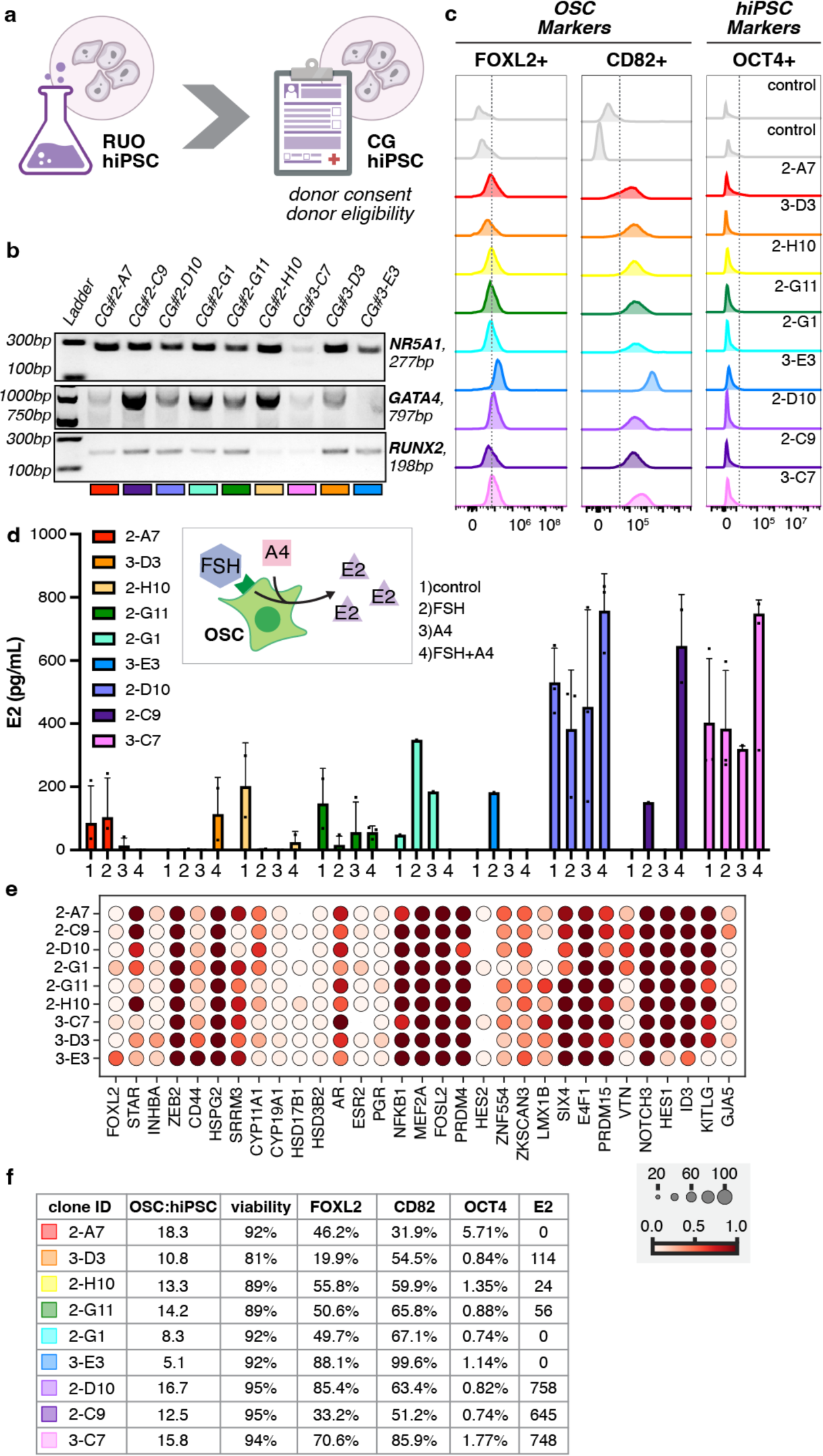
Generation and selection of a functional clinical-grade hiPSC clonal cell line as the starting material for OSC differentiation. a) Schematics of requirements for generation of a clinical-grade (CG)-hiPSC b) Image of genotyping PCR gel confirming the presence of all three transcription factors (*NR5A1*, *GATA4*, and *RUNX2*) in each of the 9 clones. c) Flow cytometry analysis of two OSC markers: FOXL2+ and CD82+, along with OCT4+ which is a hiPSC marker. The expression levels of two controls along with the 9 VCT clones were analyzed for these 3 markers. d) Quantification of estradiol (E2) levels (pg/mL) in conditioned media generated by each clone when cultured in; control media (1), control+FSH (2), control+A4 (3), and control+FSH+A4 (4), for 48 hours. Data is the mean ± SEM. e) Dotplot representing the expression of granulosa cell markers in each of the VCT-clones. Scale represents ‘Mean expression in groups’ ranging from 0 to 1, and circles represent ‘Fraction of cells in group (%) ranging from 0 to 100. f) Chart displaying the 9 clones and 4 markers: FOXL2, CD82, OCT4, and E2. The chart displays the ratio of the presence of OSC markers (FOXL2 and CD82) in relation to the hiPSC markers (OCT4 and E2) in each clone. It also displays the viability of each sample, and the amount of each marker present in each sample as a percentage.

In short, the CG-hiPSC line was engineered to harbor inducible versions of the three transcription factors known to drive differentiation into OSCs, namely *NR5A1*, *RUNX2*, and *GATA4*. Individual clones were generated by limiting the dilution of the pooled engineered population and expanded into seed banks. All the clones expanded were subjected to initial screening by genotyping PCR to confirm the integration of the three transcription factors (**Figure 5b**). Nine seed clones harboring all the transcription factors were selected to proceed with a more in-depth screening process, which included assessing their identity, potency, and safety. For that, each clone was individually differentiated into OSCs, and multiple features were recorded for screening purposes and identification of the top lead candidates. To specifically assess clonal identity, we verified expression of the OSC markers, FOXL2 and CD82 after 5 days of differentiation, and confirmed that despite the differences in the level of expression among clones, all clones were positive for both markers, suggesting overall successful generation of OSCs (**Figure 5c**). Moreover, we observed null or very low levels of the hiPSC OCT4 marker in all clones, confirming the efficacy and purity of the OSC outcome (**Figure 5c**).

To evaluate the responsiveness and functionality of individual clones, we assessed the endocrine activity of OSCs for steroidogenesis after the 5th day of differentiation. Briefly, OSCs were exposed to follicle-stimulating hormone (FSH, #2), Androstenedione (A4, #3), or a combination of both (FSH+A4, #4) hormones for 48 hours (**Figure 5d**). As previously demonstrated ^11^, functional OSCs generate estradiol (E2) in response to FSH, using A4 as a substrate (**Figure 5d**, inset). We observed that individual clones respond differently to treatment with FSH + A4 (#4), being the clones that display a greater steroidogenic response, more likely to have a better performance in maturing human oocytes (**Figure 5d**). Treatment with FSH or A4 (#2 and #3) alone allows for identification of clones that are intrinsically steroidogenic, which is an indication of an immature granulosa cell profile ^15^. To gain a more comprehensive overview of the molecular signature of the individual clones, we performed bulk RNA-sequencing of all clones individually and assessed expression of granulosa cell markers, as well as hiPSC cell markers (*POU5F1* and *NANOG*) (**Figure 5e**). All clones robustly expressed most well-known granulosa cell markers (*FOXL2*, *STAR*, *GJA5*), including genes related to important signaling pathways associated with oocyte-granulosa-cell interactions (*NOTCH3*, *HES1*, *ID3*, *KITLG*) (**Figure 5e**). These results suggested that despite the functional differences observed among clones (**Figure 5d**), differences in marker gene signatures were less pronounced among CG-OSC clones. Based on the attributes previously described, in addition to the ratio OSC:hiPSC and viability at harvest, we identified the clone 2-D10 as the top lead candidate (**Figure 5f**). This decision was informed primarily based on levels of FOXL2 and CD82 expression, as well as its responsiveness to FSH and A4 in regards to the E2 production (**Figure 5f**).

### 6. Clinical-grade hiPSC line generated for clinical manufacturing shows reproducible differentiation and comparable molecular profiling to the RUO cell line

As a way of de-risking the process by ensuring downstream safety of the selected clone, prior to transition into clinical manufacturing, we assessed and confirmed the presence of hiPSC markers on our lead candidate line, 2-D10 (hereafter referred to as CG-hiPSC), as well as confirmed cell identity and normal karyotype (**Supplementary Figure 4**). We then generated three independent CG-OSCs batches (CG-OSC-L lot 88, lot 90, and lot 116), leveraging the protocol previously identified as the most appropriate to be transitioned into clinical manufacturing. More specifically, CG-hiPSCs were differentiated with the highest grade raw material onto laminin-521 coated dishes. To evaluate the potential for scalability of the process, while the first two lots of CG-OSCs (lot 88 and lot 90) were differentiated onto 55cm^2^ dishes, as all the previous batches (Surface Unit cm^2^, **Table 1**), CG-OSC-L lot 116 was differentiated onto 175cm^2^ dishes. As expected, in all the lots cell morphology upon differentiation was characterized by small cells with granules in the cell body, tightly packed into clusters with spiky edges (**Figure 6a**). The identity of OSCs was confirmed by the expression of the markers, FOXL2 and CD82 (**Figure 6b**), and purity was verified by demonstrating that the hiPSC markers *POU5F1* (OCT4) and *NANOG* were not detected in OSCs at the end of the differentiation protocol (**Figure 6c, Supplementary Figure 5**), indicating no contamination with residual hiPSCs. We also confirmed that viability at harvest remained high, averaging 97.17±0.006% among the three batches of CG-OSCs. The ratio of OSC:hiPSC was similar to the ratio achieved with the RUO-hiPSC line when differentiated over laminin-521 (14.83±4.48), averaging at 11.41±2.19 for the first two batches (lot 88 and lot 90, **Table 1**). Interestingly, differentiation of CG-OSC-L lot 116 onto larger dishes (175cm^2^) led to considerably higher OSC:hiPSC ratio (30.7, **Table 1**), suggesting notable scalability potential.

**Figure 6:**
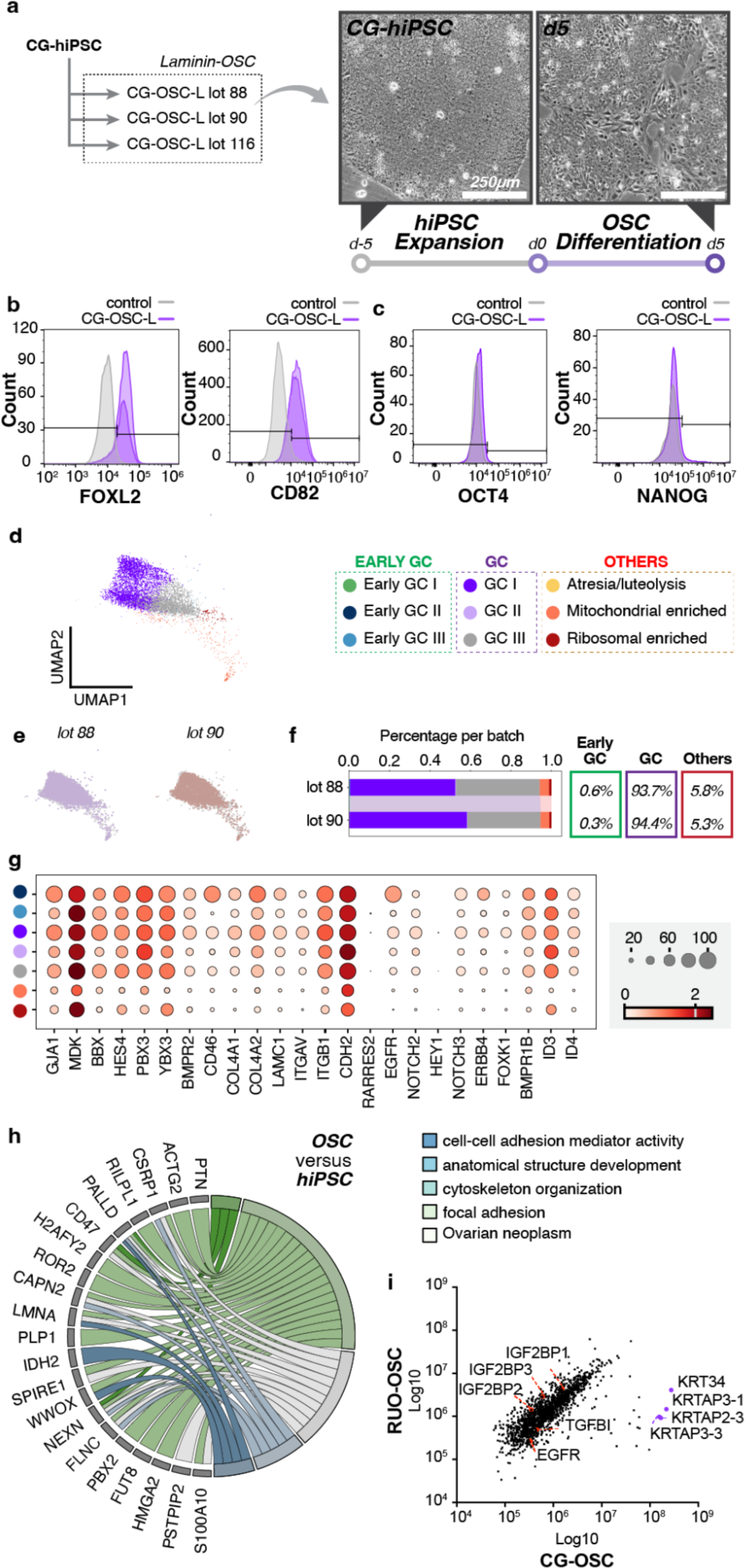
Clinical-grade hiPSC reproducibly generates a pure population of OSC. a) Images of clinical grade OSCs (lot 90) grown on laminin on day 5 of hiPSC expansion and day 5 of OSC differentiation. Scale bar, 250 μm. b) Flow cytometry analysis of 4 markers: FOXL2, CD82, OCT4, and NANOG. Expression levels of these markers were tested against a control and CG-OSC-L. d) UMAP projection of the CG-OSC subset. e) UMAP projection of the individual lots found in the CG-OSC-L subset. f) Stacked bar plot depicting the amount of each cluster type found in each lot relative to the CG-OSC-L subset. Overall percentages per group are given to the right of the barplot. g) Dotplot representing the expression of granulosa cell markers in the CG-OSC subset. Scale represents ‘Mean expression in groups’ ranging from 0 to 2, and circles represent ‘Fraction of cells in group (%) ranging from 0 to 100. h) GO chord plot for differently regulated proteins in both RUO-OSC and CG-OSC versus hiPSC. i) Correlation curve for proteins detected in the secretome of RUO-OSC versus CG-OSC.

To further characterize the transcriptional signature of the differentiated OSCs, as well as assess reproducibility among independent lots, we performed scRNA-seq in two batches of differentiated CG-OSC-L (CG-OSC-L lot 88 and 90; **Figure 6d**). Strikingly, when compared with previous samples analyzed, the two batches were nearly identical in terms of cluster distribution, and they were composed majoritarily of ‘GC’ class clusters, specially subclusters ‘GC I’ and ‘GC III’ (**Figure 6e, f**). Interestingly, the transcriptomic profile of the OSCs derived from CG-hiPSC (**Figure 6g, Supplementary Figure 6**) resembled the outstanding batch of the RUO-hiPSC line generated initially (RUO-OSC-M lot 6, **Figure 1c, f**), as well as the two batches of laminin-OSCs, generated after the raw material optimization (**Figure 3j-k**). Although not a direct measurement of the functionality of these cells, this is a good indicator that CG-hiPSC-derived OSCs will result in successful functional outcome. Additionally, this observation clearly indicates that reproducibility of final cellular outcome is consistent among independent batches, regardless of genetic backgrounds (RUO-hiPSC and CG-hiPSC lines), and even when performed by different operators. These findings are critical to ensure successful translation and to de-risk the manufacturing process towards clinical stages.

To expand our analysis beyond transcriptomics readouts, we performed proteomics of the bulk population of differentiated OSCs derived from both CG-hiPSC and RUO-hiPSC. We included in our analysis samples of undifferentiated hiPSCs from both genetic backgrounds. Despite the limited detection range of this assay compared with RNA sequencing, inclusion of these additional samples in the analysis can provide insights into the differentiation process, as well as the mechanism of action. To assess proteins and processes that were being upregulated during the differentiation process, we calculated the ratio of expression of each detected entity in OSCs at time-point 0 *versus* hiPSCs for both genetic backgrounds (**Supplementary Table 2**). Among the top 200 proteins detected with a higher ratio of expression in OSCs compared to hiPSCs, 26 were overexpressed in both cell lines and had enrichment in functional profiling terms such as “cell-cell adhesion mediator activity”, “cytoskeleton organization”, and “focal adhesion” (**Figure 6h**), suggesting that these processes are involved with OSC differentiation. Interestingly, terms related to cytoskeleton remodeling and cell adhesion were not just enriched on the shared 26 proteins by both cell lines, but also on the total top 200 proteins from each genetic background, emphasizing the importance of these processes throughout the differentiation into OSCs (**Supplementary Table 2**). Additionally, comparison of RUO-OSCs and CG-OSCs secretome has demonstrated a high correlation between these samples, further supporting comparability between both cell lines, and suggesting potential functional similarities (**Figure 6i, Supplementary Table 3**). Among the proteins secreted by both cell lines, we identified relevant players associated with oocyte maturation and developmental competency, such as TGFB1, EGFR, and IGF2BP1/2/3 (**Figure 6i, Supplementary Table 3**).

### 7. OSCs derived from clinical-grade hiPSCs consistently lead to higher rate of oocyte maturation

To further assess the comparability between CG-OSCs and RUO-OSCs in terms of functional outcomes, we cultured three independent batches of CG-OSC-L (CG-OSC-L lot 88, 90 and 116) with immature human oocytes as previously described (see Materials and Methods) and evaluated the rate of MII oocyte formation relative to the control group as the potency readout (**Figure 7a**). We observed that all the three CG-OSC-L batches successfully led to higher rates of MII maturation compared to the control (*p*=0.019, lot 88/control:1.24, lot 90/control: 1.22, lot 116/control: 1.29) in a very consistent manner (**Figure 7a**). Notably, the relative values compared to control from these three individual batches were also very similar to the relative value of RUO-OSC-L indicating that differentiation onto laminin-521 is not only reproducible among independent batches from the same cell line, but also across different genetic backgrounds (RUO-hiPSC and CG-hiPSC, **Figure 7b**).

**Figure 7:**
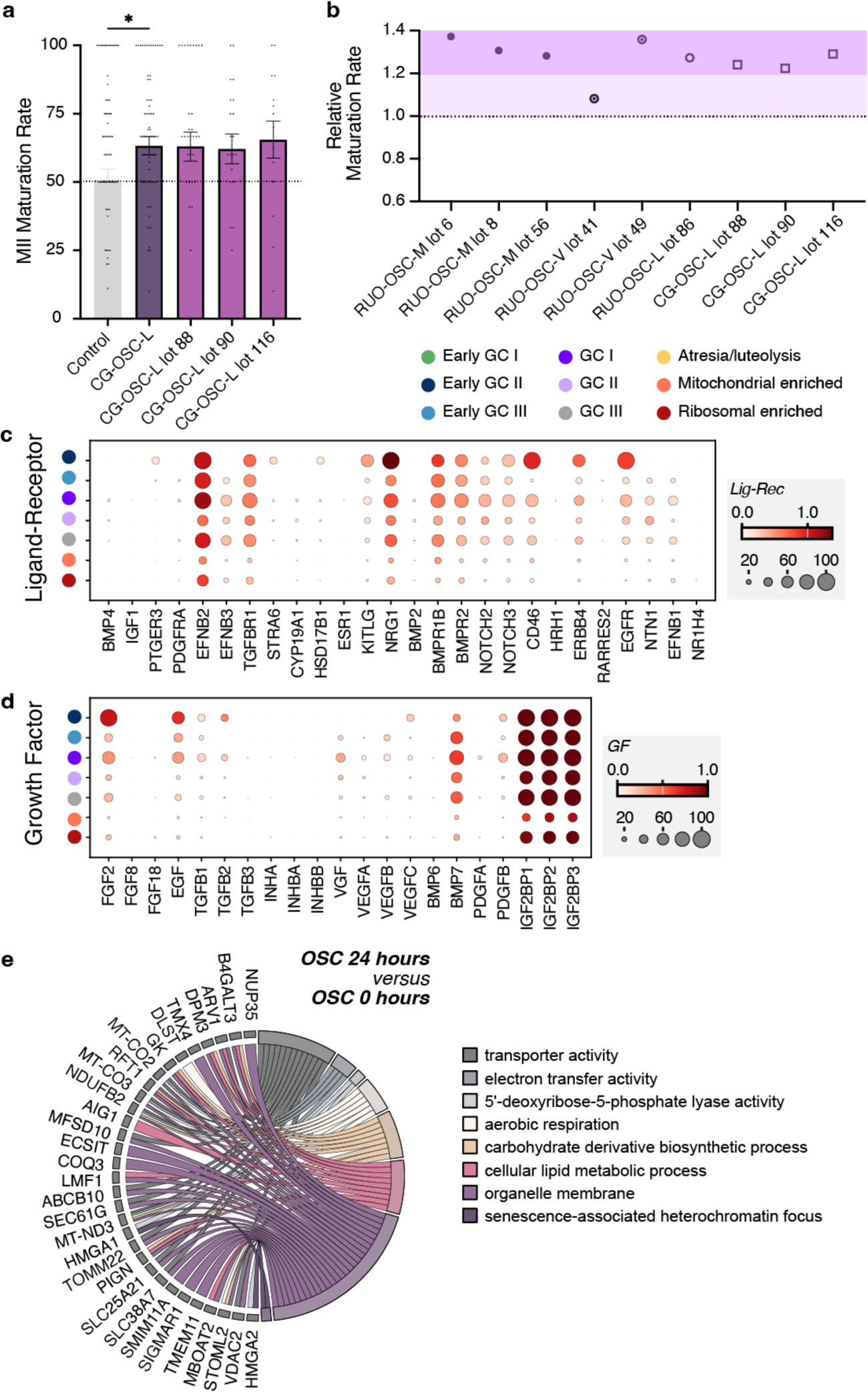
Independent batches of CG-OSC lead to robust oocyte maturation indicating successful functional outcome and comparability to RUO-OSC results. a) Comparison of MII Maturation Rates between Control-IVM group (grey) and OSC-IVM groups (CG-OSC-L, CG-OSC-L lot 88, CG-OSC-L lot 90, CG-OSC-L lot 116). Maturation rates of three separate batches (CG-OSC-L lot 88, CG-OSC-L lot 90, CG-OSC-L lot 116) are combined in the CG-OSC-L bar, with additional bars displaying the individual maturation rates of each batch. Data is the mean ± SEM (*p*=0.019, lot 88/control:1.24, lot 90/control: 1.22, lot 116/control: 1.29). b) Figure depicting the relative maturation rate per each lot, scale ranging from 0.6 to 1.4. c) Dotplot representing the expression of Ligand-Receptor Genes in the CG-OSC subset. Scale represents ‘Mean expression in groups’ ranging from 0 to 1.5, and circles represent ‘Fraction of cells in group (%) ranging from 0 to 100. d) Dotplot representing the expression of Growth Factor Genes in the CG-OSC subset. Scale represents ‘Mean expression in groups’ ranging from 0 to 1, and circles represent ‘Fraction of cells in group (%) ranging from 0 to 100. e) GO chord plots for differently regulated proteins in both RUO-OSC and CG-OSC 24 hours versus OSC at 0 hours.

Analysis of gene expression of key receptor-ligand components revealed overall enrichment of the receptors *TGFBR1*, *BMPR1B*, *BMPR2*, *NOTCH2/3*, *ERBB4*, and *EGFR*, as well as the ligands *EFNB2/3*, *NRG1*, and *NTN1* (**Figure 7c**) within the CG-OSCs batches. Notably, expression of *TGFBR1*, *BMPR2*, *NOTCH2/3*, and *EFNB2* in particular are consistently enriched in previous batches of RUO-OSCs (**Figure 2c** and **Figure 4c**), indicating their potential involvement in the OSC mechanism of action. Furthermore, growth factors identified as enriched in previous batches (**Figure 2d** and **Figure 4d**), such as *FGF2*, *TGFB1*, and *BMP7* were also enriched in the CG-OSC-L (**Figure 7d**), suggesting their pivotal role in the oocyte maturation process. This is consistent with published data demonstrating the involvement of these growth factors in orchestrating oocyte maturation through the interplay between granulosa cells and oocytes ^27–31^.

To gain insight into the potential mechanism of action of these cells during oocyte maturation beyond transcriptomics readouts, we also performed proteomics to investigate proteins that were being overexpressed in OSC after 24 hours *in vitro* in comparison with OSCs prior to culture (0 hour) from cells derived from both CG-hiPSC and RUO-hiPSC. Through a similar approach, we compared the top 200 proteins overexpressed in each genetic background and identified 40 commonly overexpressed proteins in both groups (**Supplementary Table 4**). These shared proteins were enriched for functional profiling terms such as “transporter activity”, “electron transfer activity”, “aerobic respiration”, and “cellular lipid metabolic process” (**Figure 7e**). Analysis of the top 200 proteins in each group independently underscored terms associated with metabolic processes, further suggesting the contribution of these processes for OSC function. This observation is consistent with the knowledge that glucose and lipid metabolism are fundamental metabolic pathways in granulosa cells, playing crucial roles to ensure normal oocyte development ^32^.

## Discussion

Despite historically lower efficacy overall compared to standard IVF procedures, *in vitro* maturation (IVM) has the potential to allow patients to undergo egg freezing and *in vitro* fertilization (IVF) through a safer procedure due to the significantly lower requirement for hormonal stimulation. Recently, it was demonstrated that supplementation of standard of care IVM media with OSCs leads to a higher oocyte maturation and euploid embryo formation rate ^7^, possibly due to the mechanisms employed by OSCs to recapitulate the ovarian follicle environment *in vitro* and, therefore, enhance oocyte developmental competence. Importantly, the application of this novel OSC-IVM treatment may reduce the current efficacy difference between IVM and IVF, thereby extending the applicability of this strategy beyond patients with medical need for abbreviated hormonal stimulation. This substantial improvement to IVM technology offers an alternative to current IVF practice to a wider population with significant cost savings and lower medical burden ^2^. However, translation of this technology to clinical application requires not only being able to manufacture OSCs at scale, but most importantly deep understanding of the product identity, reproducibility, and potency, while ensuring safety to the patient undergoing the procedure. Here, we showed in depth characterization of 13 batches of OSCs derived from two donor cell lines that were manufactured independently by five different operators over one year. We demonstrated across all these conditions that overexpression of the transcription factors *NR5A1*, *RUNX2*, and *GATA4* efficiently generate a pure and consistent population of OSCs (granulosa-like cells) that can range from ‘Early GC’ state to ‘atresia/luteolysis’ state, effectively covering all stages of ovarian follicle development. Optimizations applied to the manufacturing strategy do not alter this potential, but rather result in lower batch-to-batch variability in regards to cell composition, overall contributing to the product’s consistency. Moreover, we did not observe the presence of residual hiPSCs in any of the batches tested, an important safety end result for a product derived from hiPSCs ^33^. Finally, we demonstrated that by applying the same gene engineering strategy used for creating the RUO starting material (RUO-hiPSC) to a clinical-grade cell line (CG-hiPSC), differentiated OSCs maintained similar molecular, phenotypic, and functional outcomes, indicating the robustness of the process and translatability to different genetic backgrounds. The latter is an especially important finding, as it creates an opportunity to apply a similar approach for broadening our platform to encompass additional donor cell lines, including patient-derived samples, with the intention of exploring alternative applications, such as disease modeling and personalized medicine.

Applying the directed differentiation approach to drive granulosa-cell fate is crucial for achieving OSC identity efficiently and seamlessly. As previously reported by our group ^11^, the combination of transcription factors, including *NR5A1*, *RUNX2*, and *GATA4* is among the most effective approach to induce differentiation into this cell type, requiring only minimal supplementation with small molecules without the need for lengthy periods in culture. In this study, we further tested the potential synergistic effect of including small molecules and growth factors known to play a role in granulosa cell differentiation to the differentiation media, but these were shown to be unnecessary. Conversely, the substrate in which the cells are attached had a significant impact on refining molecular and phenotypic signatures of the cells within the granulosa-like spectrum, ranging from our characterization of ‘Early GCs’ to ‘GCs’. These differences in molecular signature could have a direct impact on cell behavior in long-term cultures, which in turn can impact utilization of the cells in an extended application or when specific recapitulation of morphological features is required, such as in ovarian organoids. The possibility of modulating these cellular behaviors upon selection of the substrate allows for the versatility of the platform and expands the plethora of applications.

As the mechanism of action is not yet fully understood, potency of each OSC lot has been evaluated by demonstration of MII oocyte formation rate through OSC-IVM assays. Interestingly, despite the differences observed in molecular signatures among distinct lots of OSC, functional differences were not clearly observed in this study. Notably, we have previously demonstrated that utilization of OSCs with a more immature (fetal) profile, derived from a different combination of transcription factors, is not efficient in driving human oocyte maturation when compared to the cells utilized in this study ^7^. It is still unclear whether cells in these different cellular states would be able to support efficient oocyte maturation, whether during application the cells are converted into the same potent state, or whether even a small proportion of cells would suffice to confer potency to the final product. It is also possible that analysis of a larger sample size would be necessary to evidence subtle functional differences among groups, which is a limitation due to the nature of the samples utilized for this study. The exact mechanism of action in which OSCs drive oocyte maturation is likely complex and future studies are required to fully elucidate it. It is evident that the mechanism of action relies on paracrine signaling and that direct physical interaction is not required for the cells to exert their actions. Moreover, detection of secreted growth factors by the OSCs, such as MDK, TGFB1, and IGF2BP1/2/3, which are well known to regulate oocyte maturation and developmental competence acquisition both *in vivo* and *in vitro*, further supports a potentially paracrine mechanism of action ^34–36^. We have also previously demonstrated that not only oocytes are modulated by the OSCs, but the OSC molecular signature is also modulated by the presence of oocytes ^8^. This emphasizes the importance of employing a dynamic co-culture system for IVM rather than the utilization of a static conditioned-media alternative. While the mechanism of action of OSCs during OSC-IVM is not yet fully dissected, by evaluating oocyte maturation followed by embryo formation in a surrogate species, such as mouse ^37^, informs about the therapeutic activity of the hiPSC-derived OSCs, therefore consisting of a straightforward potency assay for releasing OSC batches.

Raw material risk assessment and sourcing for alternative higher quality reagents contribute not only to reducing the risk of carrying over adventitious agents or toxins to the final product, but also to increasing reproducibility from lot-to-lot. The manner of selecting raw materials for manufacturing plays a critical role in both mitigating contamination risks and product efficacy. This is especially crucial for our application, as it involves both oocytes and ultimately IVF and live births. Furthermore, it helps to highlight and screen for process-specific concerns, such as harmful endotoxins to which oocytes are particularly sensitive. In this study, we demonstrated that this targeted approach to raw material selection fostered greater reproducibility and consistency of the final cellular outcome, which is essential for maintaining reliability and safety in clinical applications. Additionally, the process of selecting higher quality reagents highlighted a significant role of the substrate in refining the fate of OSCs, as well as increasing purity of cellular outcome and lot-to-lot reproducibility. These findings suggest that screening of different substrates may further improve and refine hiPSC differentiation processes towards multiple cell types, facilitating translation of other products towards clinical applications. In summary, the strategic selection of all raw materials serves as a cornerstone for optimizing outcomes, ensuring product quality, and maintaining the integrity of the manufacturing process.

In conclusion, this study describes generation and comparability of a clinical-grade starting material (CG-hiPSCs) to support development of a novel cellular additive for *in vitro* maturation of human oocytes (OSC-IVM), as well as manufacturing adaptations required to convert this technology into a clinical-grade product. OSC-IVM has been demonstrated to efficiently drive oocyte maturation as well as euploid blastocyst formation ^7^, and once fully integrated into clinical practice, has the potential to substantially impact patients interested in egg freezing and *in vitro* fertilization. Future challenges for the clinical manufacturing of OSC-IVM include production at scale to attend to the demand for the product (400,000-500,000 ART cycles per year in the US)^38^, which will require automation and system closure to streamline and de-risk the process. Despite these challenges, we strongly believe that this technology will constitute not only the starting material for OSC-IVM products, but also serve as a toolkit with the potential to be further expanded into multiple applications and indications focused on women’s health and infertility.

## Acknowledgement

This work was performed with the support of clinical partnerships at Spring Fertility New York, Extend Fertility, Ruber Clinic of Madrid, Tambre Clinic of Madrid, and Pranor Clinic of Lima. We thank the dedicated support and work of the embryology and support staff teams at these clinics for coordinating and managing the collaborative study, as well as the technical support of Christopher Collado and Kariana Flores. We additionally thank the Wyss Institute for Biologically Inspired Engineering at Harvard University for material transfer of reagents used in this preclinical work. We thank Professor Mary Herbert, Professor Phillip Jordan, Professor George Church, Professor Kristin Baldwin, Dr Sara Vaughn, and Professor David Albertini for advice and guidance on the use of OSCs in IVM work. We also thank the New York University flow cytometry and imaging cores, the Proteomics and Metabolomics Core Facility at Weill Cornell Medicine, and Azenta/Genewiz for their assistance in data generation and analysis.

## Author Contribution

C.C.K., B.P., F.B., A.D.N, and S.P. conceived the experiments. S.P., M.M., A.G., and F.B. performed all embryology work for the study, C.C.K. supervised their work. B.P., M.J., K.S.P., A.B.F., A.D.N., and F.B. produced and qualified OSC batches, C.C.K. supervised their work. B.P., S.K., and G.R. worked on bulk- and scRNA-seq analysis and cell type assignments. F.B. performed proteomic experiments and analysis. A.B.F. coordinated the assay logistics. C.C.K., B.P., F.B., and A.D.N. wrote the manuscript with significant input from all authors.

## Methods

### Source material

The research-use-only hiPSC (RUO-hiPSC) was sourced from the laboratory of G. Church ^11^. A female donor hiPSC line (VCT-37-F35) was sourced from Reprocell USA (9000 Virginia Manor Rd #207, Beltsville, MD 20705) to serve as the starting material for our clinical-grade cell line. The stem-cell line, derived from human-skin fibroblasts, was generated under GMP conditions using a non-integrative, mRNA-based reprogramming technology with controlled conditions and GMP-compliant reagents used for the entirety of the manufacturing process. Regarding specific stages in that process, proper controls were implemented for fibroblast derivation according to established guidelines, while reprogramming and cell expansion took place under fully GMP conditions in compliance with regulatory standards and guidelines of the FDA, EMA, and PMDA. Donor eligibility was determined to be in accordance with 21 CFR Part 1271 Subpart C and FDA Guidance for Industry: Eligibility Determination for Donors of Human Cells, Tissues, and Cellular and Tissue Based Products (HCT/Ps), 2021. All of the experiments involving human cells were performed according to ISSCR 2021 guidelines (add reference), and approved by the IRB committees.

### Plasmid Manufacturing

Plasmids utilized for engineering were manufactured as previously described ^11^. Whole plasmid sequencing was performed by Plasmidsaurus using Oxford Nanopore Technology with custom analysis and annotation. All plasmids were further screened for purity and stored at -20°C, while glycerol stocks of transformed bacteria are stored at -80°C.

### Cell Engineering

Engineering of hiPSCs with specific transcription factors, including *NR5A1*, *RUNX2*, and *GATA4*, was performed using a piggyBac transposase strategy and the Lonza Nucleofector system, as previously described ^11^. Puromycin (Sigma-Millipore) selection was utilized to eliminate cells without integration of the transcription factors. Insertion sites into the CG-hiPSC line were verifiable by Whole Genome Sequencing (Azenta). From the analysis, there was also no evidence of mutations on 20 commonly known proto-oncogenes.

### Cell screening, selection, and preliminary characterization

Following the preliminary round of testing on the pooled population of transfected hiPSCs, clones were established by limiting dilution in multiwell plate format. All wells were closely monitored daily until identification of single clones in each well. Wells with more than one clone identified were discarded. Each identified clone was further expanded and cryopreserved. Each clone was initially assigned a unique code. All seed clones were subjected to genotyping PCR to identify the presence of the three transcription factors. This initial screening resulted in identification of nine clones harboring all the transcription factors, each of which was then subjected to a more in-depth screening process, including identity, potency, and safety assays for the identification of a lead candidate cell line. To identify the lead candidate, each of the nine clones was individually differentiated and subjected to a series of assays to ensure identity (pluripotency markers, and genotyping), conformance (cell count and viability), and potency (OSC production and function) of the clones. The leading candidate clone (2-D10) selected to be used as the starting material for the clinical-grade cell line was named CG-hiPSC.

### hiPSC Maintenance and OSC Differentiation

hiPSC were maintained in feeder-free conditions and cell culture plates were pre-coated with either Matrigel (Corning), Vitronectin-XF (StemCell Technologies), Laminin-521 (StemCell Technologies), or Laminin-511 (Reprocell). Cells were maintained in either mTESR1 (StemCell Technologies) or TeSR-AOF (StemCell Technologies) without antibiotics, at 37 °C in 5% CO_2_. All human pluripotent stem cells were negative for mycoplasma or other human adventitious agents (performed by IDEXX Bioanalytics), and karyotypically normal (G-band karyotype test, performed by WiCell Research Institute, and Karyostat, performed by ThermoFisher). RUO-hiPSC and CG-hiPSC were authenticated by SNP array (CellID, performed by ThermoFisher). OSC differentiation was performed as previously described ^11^. In short, hiPSCs were exposed to the Rho kinase inhibitor, Y-27632, and the WNT activator, CHIR99021, to prime the cells into a mesodermal fate. Exposure to doxycycline throughout the entire process induces overexpression of the transcription factors, *NR5A1*, *RUNX2*, and *GATA4*, directing differentiation of hiPSCs towards OSCs. The differentiation process required five days in culture. All images were taken with ECHO Revolve Microscope (Discover ECHO).

### Cell Count and Viability

Cell counts and viability assessment was performed using Eve Automated Cell Counter (NanoEnTeck) and NucleoCounter NC-202 (ChemoMetec).

### Single cell RNA Sequencing (scRNA-seq)

OSCs were cryopreserved in CryoStor CS10 (StemCell Technologies) prior to being processed for single-cell RNA sequencing by Genewiz (Azenta). Cells were loaded onto a Chromium Single Cell Chip (10x Genomics), and processed through the Chromium Controller to generate single-cell gel beads in emulsion. scRNA-seq libraries were generated using the Chromium Single Cell 3′ Library & Gel Bead Kit (10x Genomics). Target cell recovery was estimated to be 3,000 cells per sample. Files were generated following a split-set analysis workflow. First bcl2fastq was used to generate the fastq files. Next, a reference genome was generated using split-pipe and the Homo sapiens GRCh38 file, also using STAR^39^ and Samtools^40^. Then, split-pipe was run to process and align the files against the reference genome. The resulting files were generated in an mtx matrix format. Finally, the files were combined, and an Anndata object was made in h5ad file format. For the analysis, cells with less than 200 genes were filtered out, as well as genes that were found in less than 3 cells. Cell counts were normalized to 10,000 UMIs (Unique molecular identifiers) per sample and log (ln) plus 1 transformed. Principal component analysis was performed using the Scanpy package (v1.9.6) based on 30 PCA components, and using PCA results, nearest neighbor analysis was performed using n_neighbors = 20. Batch correction was performed using Scanpy’s ComBat method^41^. Number of components used for batch correction was also 30 and the data was then transformed using the Uniform Manifold Approximation and Projection (UMAP) method^42^. Clusters were formed using the Leiden method^43^ with a resolution of 0.25 for the complete dataset (containing all lots described in the manuscript), and projected into the subsetted objects. At first, 15 Leiden clusters were identified. Thereafter, clusters were combined based on biological similarities, which resulted in nine final clusters: Early GC I, Early GC II, Early GC III, GC I, GC II, GC III, Atresia/luteolysis, Mitochondrial gene enriched (Atresia/luteolysis), and Ribosomal gene enriched (Atresia/luteolysis), (**Supplementary Figure 7, Supplementary Table 5**). The marker genes per cluster, as well as certain granulosa cell markers (*GJA1, MDK, BBX, HES4, PBX3, YBX3, BMPR2, CD46, COL4A1, COL4A2, LAMC1, ITGAV, ITGB1*), were analyzed in order to identify cluster cell types (**Supplementary Table 5**). Dot plots and feature plots were generated to assess the expression levels of certain genes either per cluster or per sample. Based on downstream analysis, clusters 0 and 5 were subsetted from the original object and they were re-clustered using the Leiden method at a higher resolution of 0.3. Subclusters were combined on the basis of biological similarities. Once all the subgroups were identified, the subclusters resulting from cluster 0 and cluster 5 were merged together with the original object. Gene signatures based on genes in the folliculogenesis stage ^14^ were also analyzed and used in predicting cluster identification. Signature scores for the RUO-OSC-M subset were generated for two groups: Antral and Pre-Ovulatory genes (**Figure 1f)**. Based on the final object, 4 more subsets were generated: (1) a RUO-OSC-M object consisting of the following lots: lot 6, lot 7, lot 8, lot 29, lot 48, and lot 56; (2) a RUO-OSC-V object consisting of the following lots: lot 41, lot 49, and lot 57; (3) a RUO-OSC-L object consisting of lot 77 and lot 86; and (4) a CG-OSC-L object consisting of lot 88 and lot 90. Some samples were excluded from the subsets due to low-scale OSC production, includingRUO-OSC-V lot 37, RUO-OSC-V lot 39 and CG-OSC-V lot 0. UMAPS of all the subsets were generated with the cluster names from the original object (**Figures 1c, 3e, 3f, 6d**). Individual UMAPS of each lot were also generated (**Figures 1c, 3h, 3j, 6e**). Lists of markers from groups like: granulosa cells genes, pre GC I/II genes, and steroidogenesis related genes (E2) were made for further analysis. A dot plot for the RUO-OSC-M subset was made depicting the expression of all these groups of genes (**Figure 1d-e**). Dot plots of the granulosa cell genes (GC genes) were also generated for the RUO-OSC-V, RUO-OSC-L, and CG-OSC subsets (**Figures 3g and 7g**). Additional dot plots were also generated for all subsets with the following groups: Ligand-receptor genes and growth factor-related genes (**Figures 2c, 2d, 4c, 4d, 7c, 7d**). Stacked bar plots indicating the percentage of each cell type in each sample were created in order to validate consistency and similarity amongst the samples (**Figures 1g, 3i, 3k, 6f**).

### Bulk RNA-sequencing

Libraries for RNA sequencing were generated using the NEBNext Ultra II Directional RNA Library Prep Kit for Illumina (NEB #7765L) in conjunction with NEBNext Multiplex Oligos for Illumina (Unique Dual Index UMI Adaptors RNA Set1, NEB #7416S) and NEBNext Poly(A) mRNA Magnetic Isolation Module (NEB #E7490L), according to the manufacturer’s instructions. Library pool was sequenced at Azenta using Illumina 2x150bp, ∼350M PE reads (∼105GB), lightening package. Illumina sequencing files (bcl-files) were converted into fastq read files using Illumina bcl2fastq (v2.20) software deployed through BaseSpace using standard parameters. RNA-seq data gene transcript counts were aligned to *Homo sapiens* GRCH38 (v2.7.4a) genome using STAR (v2.7.10a) ^39^ to generate gene count files and annotated using ENSEMBL ^39^. Gene counts were combined into sample gene matrix files (h5). Computational analysis was performed using the Scanpy (v1.9.6) package. Two h5ad files were joined on the basis of similar features and genes. The two merged files were created into one Anndata object which was normalized to 10,000 UMI per sample and log (ln) plus 1 transformed. Principal component analysis was performed using 30 PCA components. Projection into two dimensions was performed using the Uniform Manifold Approximation and Projection (UMAP) method ^42^. This analysis was performed to evaluate the CG-hiPSC sub-clones: 2-A7, 2-C9, 2-D10, 2-G1, 2-G11, 2-H10, 3-C7, 3-D3, and 3-E3. A dot plot using Scanpy’s software, containing granulosa cell markers, was generated for all 9 clones to demonstrate gene expression in each (**Figure 5e**).

### Design of Experiments (DOE)

Design of Experiments (DOE) was conducted using JMP software (JMP 17, SAS Institute Inc., Cary, NC, USA) to optimize experimental parameters, including media supplementation, substrates, and doxycycline treatment designs, for OSC differentiation. Factors were selected from established modulators for OSC specification in the literature and factor ranges were determined through literature review (**Supplementary Table 1**). Responses were FOXL2 expression and viability. Viability was used to screen and remove experimental groups below 50% cell survival, which would be unworkable in a manufacturing context, prior to the final statistical analysis. A custom design including center points was employed and optimized for D-optimality to investigate main effects and interactions. The JMP software facilitated generation of the experimental design matrix, as well as statistical analysis through response surface methodology (RSM) and ANOVA and optimization of conditions to maximize response desirability for FOXL2 expression, allowing for exploration of the parameter space and identifying optimal conditions and primary drivers of the targeted OSC state.

### Proteomics

Liquid chromatography followed by tandem mass spectrometry (LC-MS/MS) analysis was conducted on a series of samples, including RUO-hiPSC (n=1), CG-hiPSC (n=1), RUO-OSC-M lot 56 at time 0h (n=1) and after 24h (n=1) of culture with supplemented MediCult IVM media, and CG-OSC-L lot 88 at time 0h (n=1) and after 24h (n=1) of culture with supplemented MediCult IVM media. Each condition involved the analysis of 2 million cells. Supplemented IVM media consisted of MediCult IVM media (Origio) supplemented with 75 mIU/mL of recombinant FSH (Millipore), 100 mIU/mL recombinant hCG (Sigma), 500 ng/mL of androstenedione (Sigma), 1 ug/mL of doxycycline (StemCell Tech) and 10 mg/mL of human serum albumin (HSA; Life Global). Conditioned media derived from RUO-OSC-M lot 56 (n=1) and CG-OSC-L lot 88 (n=1) was also analyzed using LC-MS/MS. To generate conditioned media, 2 million OSC cells were cultured in 2 ml of supplemented MediCult IVM media for 24 hours, maintaining the ratio of the intended clinical cell dose of 1,000 OSC cells per 1µl of media. The 24-hour culture was performed on an incubator with CO_2_ set for a pH of 7.2-7.4. Following culture, OSC cells and conditioned media were separated and processed independently. Supplemented IVM media without OSCs was used as a media control. The conditioned media were subjected to consecutive centrifugations (300g, 1,200g, and 3,000g) to remove cellular remnants, and then passed through albumin depletion columns (AVK-50, AlbuVoid Albumin Depletion Columns Biotech Support Group) to eliminate HSA-derived albumin. Proteins from both cells and conditioned media were precipitated using acetone, re-suspended in 0.1% RapiGest and 25 mM ammonium bicarbonate, reduced with DTT, and alkylated with iodoacetamide, before undergoing in-solution trypsin digestion overnight at 37dC. The resulting peptides were desalted using C18 stage-tip columns prior to analysis using a Thermo Fisher Scientific EASY-nLC 1200 coupled online to a Fusion Lumos mass spectrometer (Thermo Fisher Scientific). Buffer A (0.1% FA in water) and buffer B (0.1% FA in 80 % ACN) were used as mobile phases for gradient separation. For peptide separation, a packed in-house 75 µm x 15 cm chromatography column (ReproSil-Pur C18-AQ, 3 µm, Dr. Maisch GmbH, German) was used. Peptides were separated with a gradient of 5–40% buffer B over 30 min, and 40%-100% B over 10 min at a flow rate of 400 nL/min. Fusion Lumos mass spectrometer operated in a data independent acquisition (DIA) mode, collecting MS1 scans in the Orbitrap mass analyzer from 350-1400 m/z at 120K resolutions. The instrument was set to select precursors in 45 x 14 m/z wide windows with 1 m/z overlap from 350-975 m/z for HCD fragmentation. MS/MS scans were collected in the orbitrap at 15K resolution. Data analysis involved searching against human Uniprot database (8/7/2021) using DIA-NN v1.8 with filtering for 1% false discovery rate (FDR) for both protein and peptide identifications. Protein intensities were normalized and log transformed for relative quantitation, and multiple hypothesis correction of p-values was performed using the Benjamini-Hochberg method. Proteomic analyses were conducted at the Proteomics and Metabolomics Core Facility at Weill Cornell Medicine (New York, USA). GO Chord graphs generated with the free online platform, SRplot ^44^.

### Immunostaining (Immunofluorescence and Flow Cytometry)

Immunofluorescence staining was conducted on fixed hiPSCs adhered to a slide, following the protocol recommendations from the New York Stem Cell Foundation (NYSCF). Briefly, hiPSCs were fixed in 4% paraformaldehyde (PFA), followed by a blocking step in a blocking buffer (3% donkey normal serum and 0.1% Triton-X). The primary antibodies used were mouse monoclonal antibody against OCT3/4 (1:200; sc5279, Santa Cruz Biotechnology), goat polyclonal antibody against SOX2 (1:50; AF2018, R&D systems), goat polyclonal antibody against NANOG (1:50; AF1997, R&D systems), and Alexa Fluor 488 mouse monoclonal antibody against TRA-1-60 (1:100; 560173, BD Biosciences). The secondary antibodies used were Alexa Fluor 555 donkey anti-mouse IgG (A32773, Invitrogen), Alexa Fluor 488 donkey anti-goat IgG (A32814, Invitrogen), and Alexa Fluor 647 donkey anti-goat IgG (A32849, Invitrogen). All antibody dilutions were prepared in a blocking buffer and incubated at room temperature (RT) for 1 hour. After incubations, samples underwent three washes of 30 min each with PBS containing 0.1% Tween-20 (PBST). Subsequently, samples were mounted with Prolong Gold mounting medium prior to imaging using an ECHO Revolve microscope.

Flow cytometry analyses were conducted on RUO-OSC and CG-OSCs. For the analysis of live cells, cells were incubated with a PE-conjugated mouse monoclonal antibody against CD82 (1:50 dilution; 342104, BioLegend) in FACS wash (dPBS with 5% fetal bovine serum (FBS)). After incubation, cells were washed with FACS wash, stained with propidium iodide (1:20 dilution; P4864, Millipore Sigma) for live/dead cell staining, and subsequently analyzed using a CytoFlex Flow Cytometer. For the analysis of fixed cells, cells were fixed with 4% PFA for 15 minutes at RT and then washed with dPBS. After, cells were permeabilized using FACS wash solution containing 0.1% Triton X-100 (A16046.AE, Thermo Fisher Scientific). The primary antibodies used were mouse monoclonal antibody against OCT3/4 (1:50 dilution; sc5279, Santa Cruz Biotechnology), and rabbit polyclonal antibody against FOXL2 (1:100 dilution; A16244, ABclonal). The secondary antibodies used were Alexa Fluor 555 donkey anti-mouse IgG (A32773, Invitrogen), and Alexa Fluor 488 donkey anti-rabbit IgG (A32790, Invitrogen). Following incubations, cells were washed with FACS wash containing Triton X-100, and then analyzed using a CytoFlex Flow Cytometer. Unstained cells (negative controls) were used to determine the gating strategy.

### RT-qPCR and Genotyping PCR

For genotyping PCR, DNA extraction from various hiPSC clones was carried out using the QuickExtract DNA Extract Solution (Epicentre), following the manufacturer’s instructions. PCR amplification was performed using Q5 High-Fidelity 2X Master Mix (New England Biolabs) for 35 cycles with a 20-second extension time. Subsequently, PCR was performed to validate the integration of the 3 transcription factors *NR5A1*, *GATA4*, and *RUNX2*. The PCR protocol involved an initial denaturation step at 98°C for 30 seconds, followed by 35 cycles of denaturation at 98°C for 10 seconds, annealing at 66-70°C (*NR5A1*, 67°C; *RUNX2*, 66°C; *GATA4*, 70°C) for 10 seconds, and extension at 72°C for 20 seconds, with a final extension step of 2 minutes. The reaction was then held at 4°C. Gel electrophoresis (2% agarose) was performed to confirm the presence of insertions.

RT-qPCR was performed to assess gene expression markers of *POU5F1* and *NANOG*, following the protocol recommendations from the New York Stem Cell Foundation (NYSCF). RNA extraction was performed using the Quick-RNA Microprep Kit (Zymo Research) following the manufacturer’s instructions. cDNA synthesis was carried out with the LunaScript RT SuperMix Kit (New England Biolabs), using a thermocycler program consisting of a primer annealing stage at 25°C for 2 minutes, followed by cDNA synthesis at 55°C for 10 minutes, concluded with heat inactivation at 95°C for 1 minute. Quantification of RNA and cDNA was performed using Nanodrop. PowerUp SYBR Green Master Mix (Applied Biosystems) was used for RT-qPCR. The RT-qPCR protocol involved an initial denaturation step at 95°C for 2 minutes, followed by 40 cycles of denaturation at 95°C for 3 seconds, annealing at 60°C for 30 seconds, and an analysis step.

### Functional Assessment (Oocyte maturation)

The oocyte maturation-stimulating potential of various OSC batches was used to evaluate the potency of each batch. Briefly, immature oocytes surrounded by cumulus cells, known as cumulus-oocyte complexes (COCs), were retrieved from subjects that underwent minimal stimulation of hormonal protocols. Subsequently, these immature COCs were co-cultured with different batches of OSC cells for 24-30 hours to facilitate *in vitro* maturation (IVM). After IVM, oocytes were evaluated for their maturation state and categorized into immature stages (GV oocyte or MI oocyte) or mature stage (MII oocytes). MII oocyte maturation rate (%) was calculated by dividing the total number of mature MII oocytes by the initial number of immature oocytes, and used as the potency readout. For a comprehensive description of the methods follow, refer to our recent publication^7^. Sibling oocytes were used for initial comparisons described in **Figure 2b**. For all the other comparisons, control group and OSC-IVM group consist mostly of oocytes from non-overlapping donor cohorts. All analyses were performed using One-way ANOVA (Dunnett’s multiple comparison test). The number of oocyte donor participants and the total number of oocytes used per OSC batch assessed are detailed in **Supplementary Table 6**.

### Subject ages, ethics, and informed consent

This study was performed according to the ethical guidelines outlined in the Declaration of Helsinki. Oocyte donor participants were enrolled in the study at several fertility clinics, including the Ruber Clinic (Madrid, Spain), Spring Fertility Clinic (New York, USA), Extend Fertility Clinic (New York, USA), and Pranor Clinic (Lima, Peru), using informed consent for donation of gametes for research purposes, with ethical approval from CNRHA 47/428973.9/22 (Spain), Western IRB No. 20225832 (USA), and Protocol No. GC-MSP-01 (Peru), respectively.

## Supplementary Figures

**Supplementary Figure 1:**
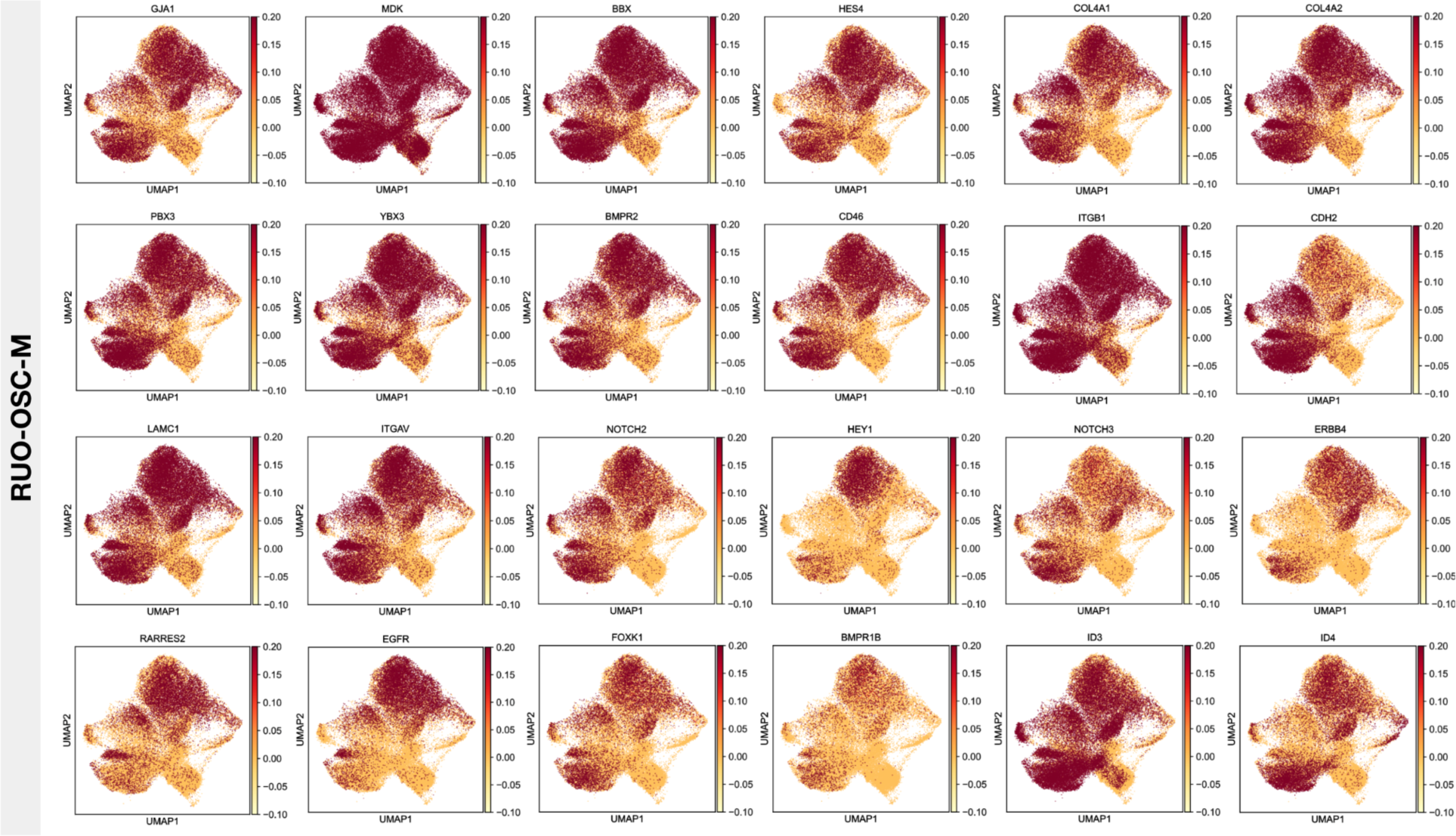
Distribution of granulosa cell markers within research-use-only (RUO) ovarian support cell (OSC) population. Feature plots depicting expression of granulosa cell marker genes across the RUO-OSC-M subset. M: matrigel. Referent to Figure 1d.

**Supplementary Figure 2:**
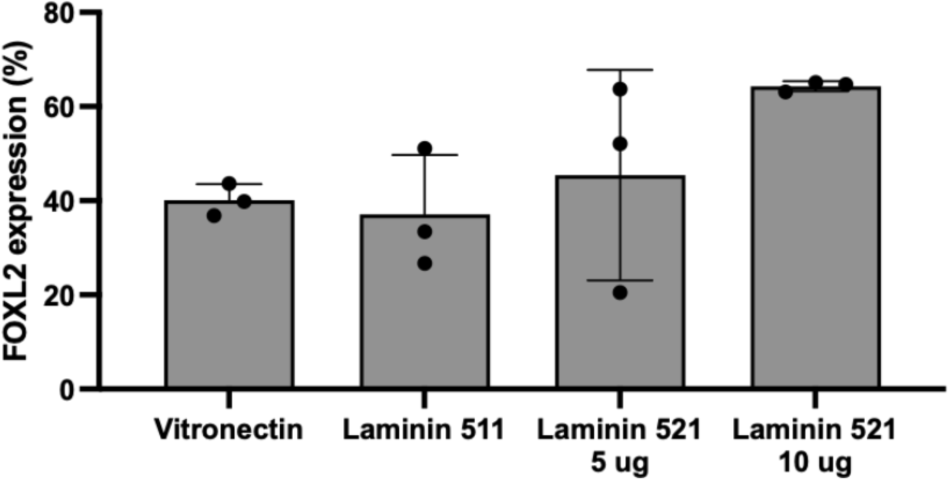
Expression of FOXL2 across different substrates. Bar plot demonstrating percentage of FOXL2 expression measured by flow cytometry on day 5 of OSC differentiation onto multiple substrates. Referent to Figure 3a-b.

**Supplementary Figure 3:**
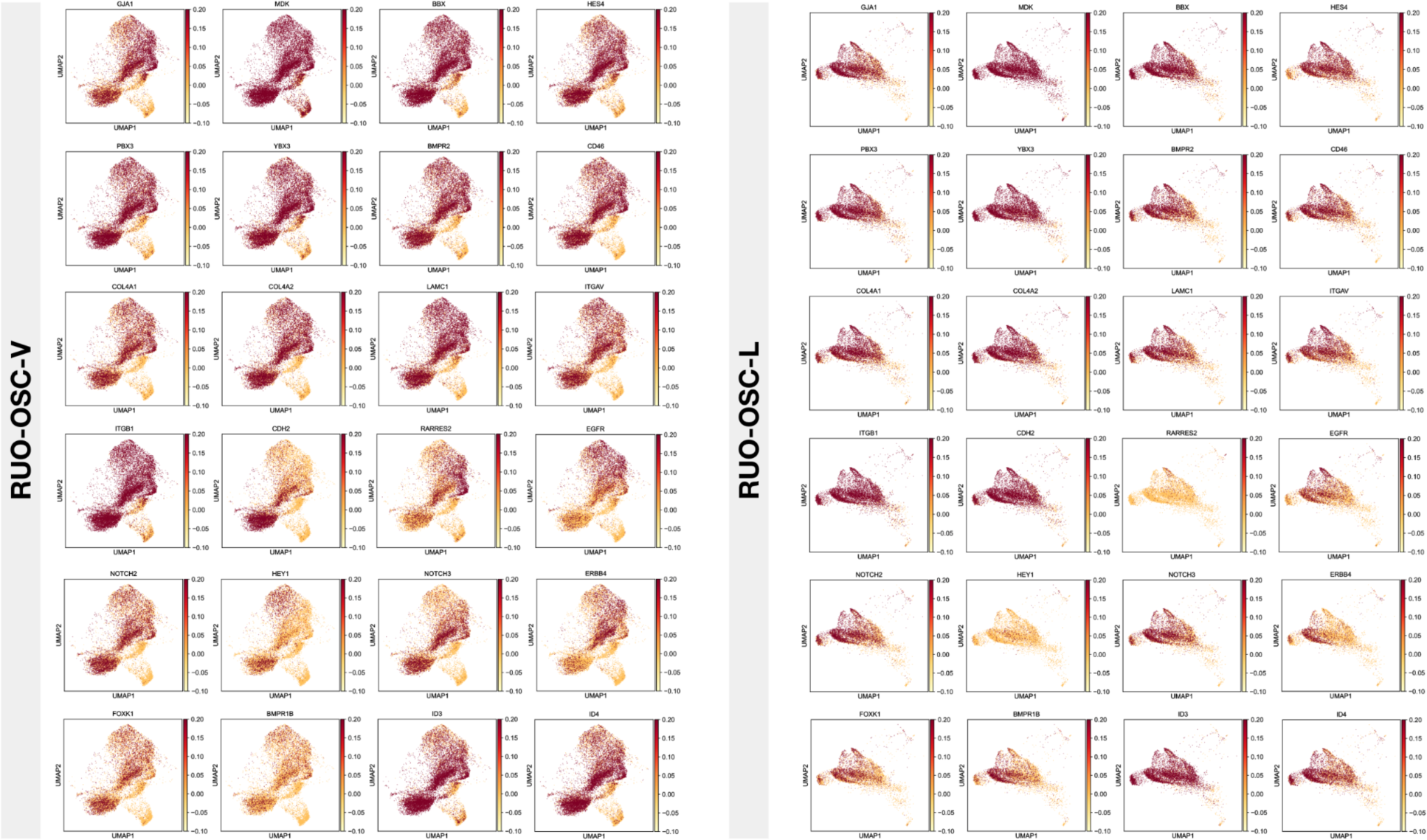
Distribution of granulosa cell markers within research-use-only (RUO) ovarian support cell (OSC) population differentiated in xeno-free conditions. Feature plots depicting expression of granulosa cell marker genes across the RUO-OSC-V/L subsets. V: Vitronectin, L: Laminin. Referent to Figure 3g.

**Supplementary Figure 4:**
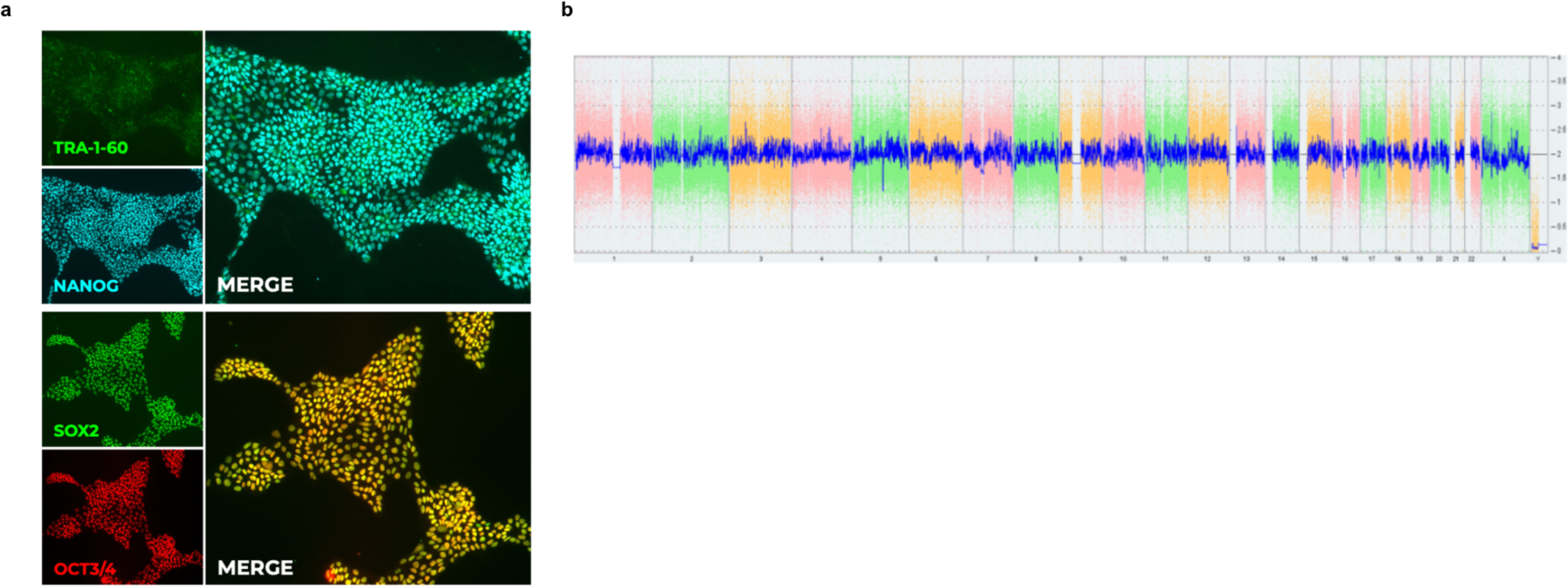
Characterization of engineered clinical-grade (CG) human induced pluripotent stem cell (hiPSC) lead clone. a) Immunocytochemistry for the hiPSC related markers, TRA-1-60 (upper panel, green), NANOG (upper panel, blue), SOX2 (lower panel, green), and OCT3/4 (lower panel, red) performed on expanded clone. b) Karyostat assay (ThermoFisher) demonstrating no abnormalities detected. The whole genome view displays all somatic and sex chromosomes in one frame with high level copy number. The smooth signal plot (right y-axis) is the smoothing of the log2 ratios which depict the signal intensities of probes on the microarray. A value of 2 represents a normal copy number state (CN = 2). A value of 3 represents chromosomal gain (CN = 3). A value of 1 represents a chromosomal loss (CN = 1). The pink, green and yellow colors indicate the raw signal for each individual chromosome probe, while the blue signal represents the normalized probe signal which is used to identify copy number. Referent to Figure 6a.

**Supplementary Figure 5:**
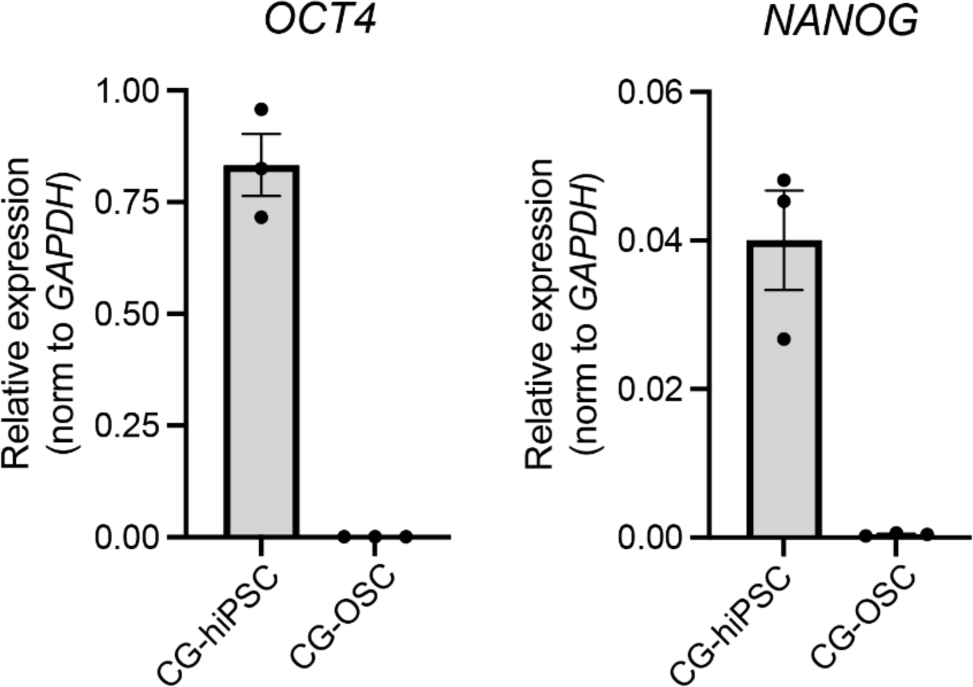
Residual human induced pluripotent stem cells (hiPSC) are not detected in the differentiated population of ovarian support cells (OSCs). Bar plots depicting relative expression of hiPSC related markers, OCT4 and NANOG in OSCs after 5 days of differentiation (CG-OSCs). Clinical-grade (CG)-hiPSC are used as a positive control. Expression is normalized by expression of the housekeeping gene, GAPDH. Referent to Figure 6b.

**Supplementary Figure 6:**
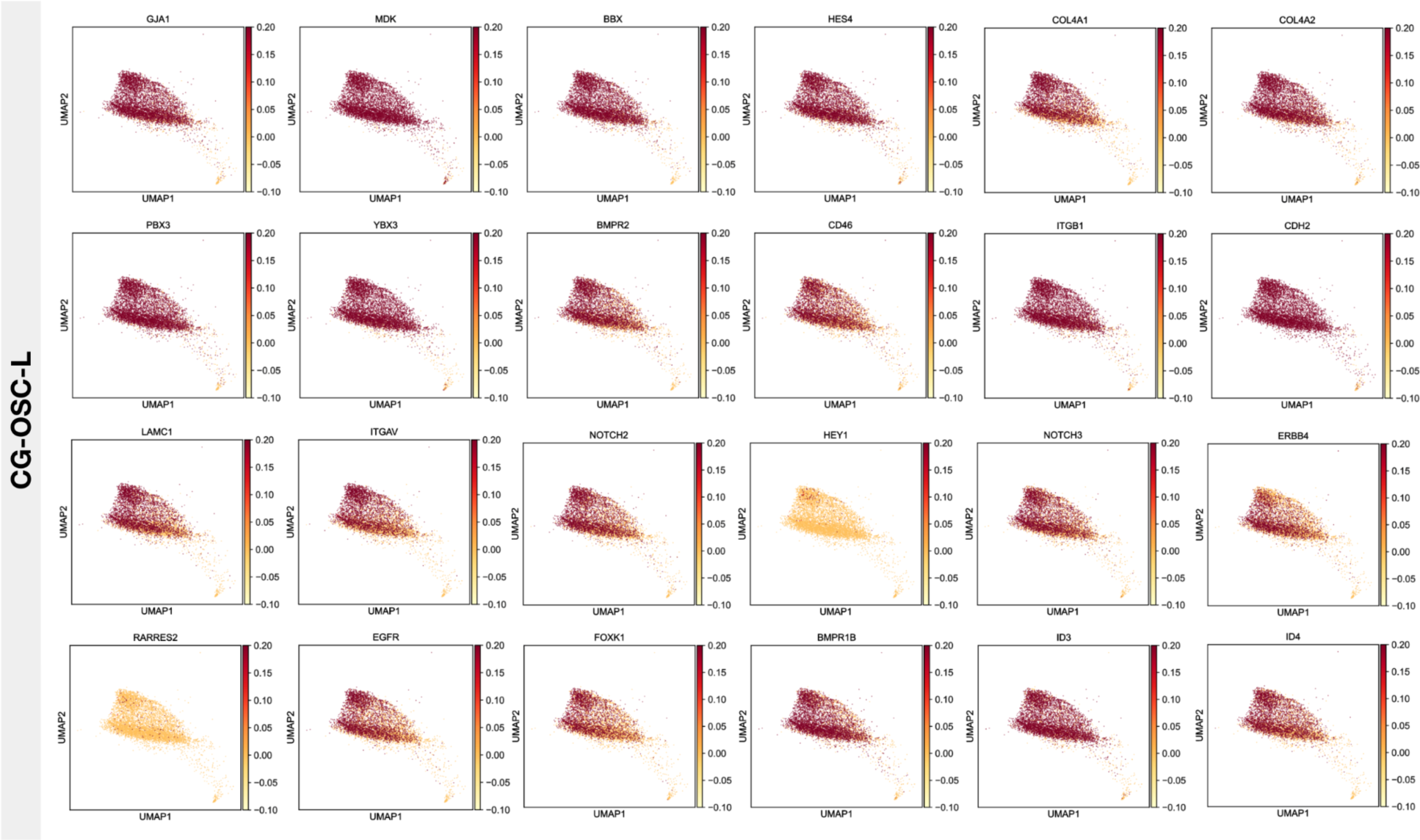
Distribution of granulosa cell markers within clinical-grade (CG) ovarian support cell (OSC) population. Feature plots depicting expression of granulosa cell marker genes across the CG-OSC-L subset. L: Laminin. Referent to Figure 6g.

**Supplementary Figure 7:**
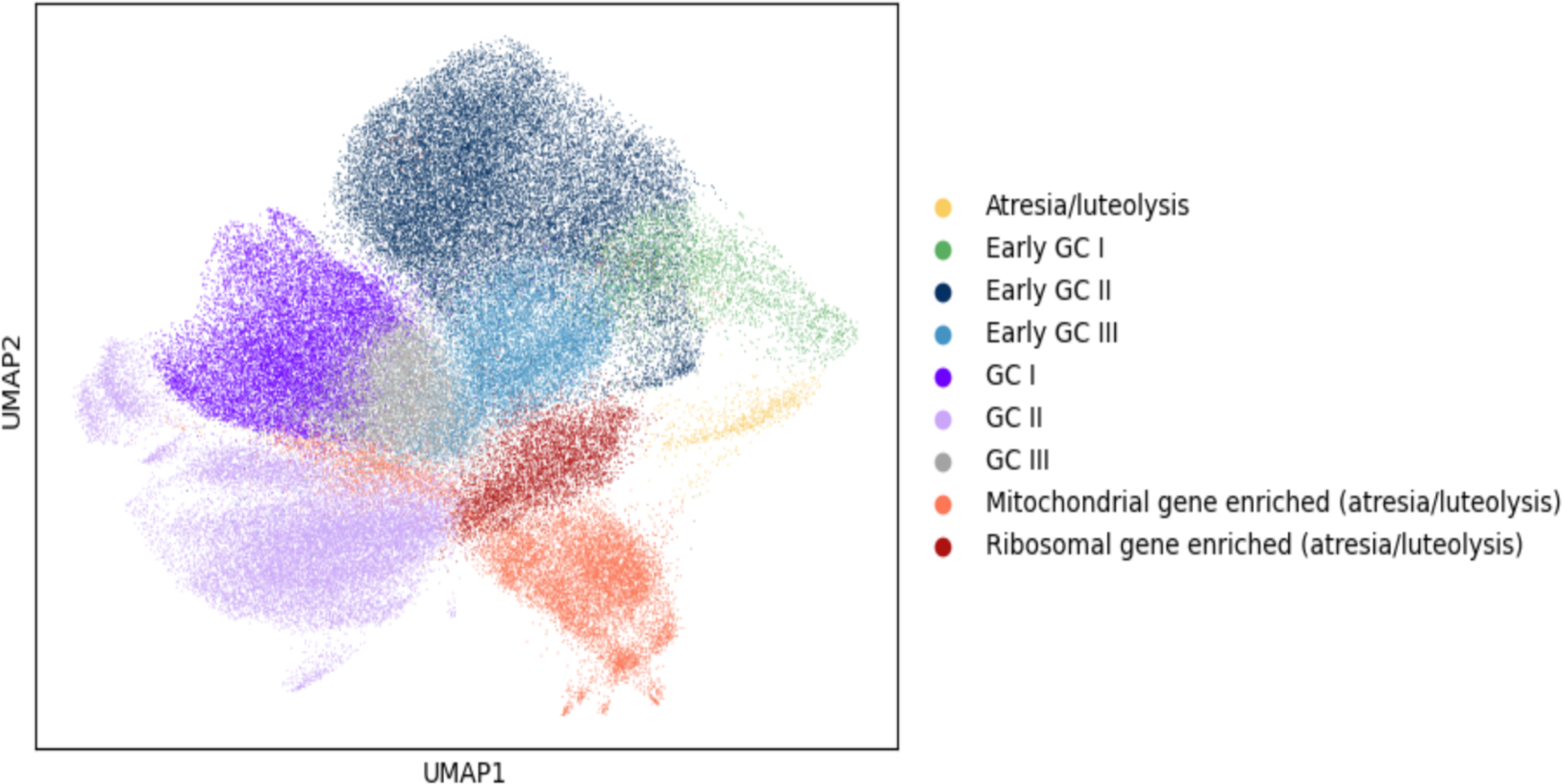
Uniform Manifold Approximation and Projection (UMAP) of the entire ovarian support cell (OSC) dataset. UMAP encompasses the following subsets: RUO-OSC-M, RUO-OSC-V, RUO-OSC-L, and CG-OSC-L. RUO: Research-use-only, M: Matrigel, V: Vitronectin, L: Laminin, CG: Clinical-grade.

